# High-fat diet exacerbates the loss of bone fracture toughness in aging for C57BL/6JN mice

**DOI:** 10.1101/2024.08.13.607633

**Authors:** Kenna Brown, Ghazal Vahidi, Brady D. Hislop, Maya Moody, Steven Watson, Hope D. Welhaven, Ramina Behzad, Kat O. Paton, Lamya Karim, Ronald K. June, Stephen A. Martin, Chelsea M. Heveran

## Abstract

The elderly are at increased risk of bone fracture and more often consume poor quality diets, such as high-fat diet (HFD). We hypothesized that HFD would exacerbate the loss of bone fracture resistance in aging. To test this hypothesis, we fed 5-month and 22-month C57BL/6JN mice of both sexes HFD or low-fat diet for 8 weeks. Aging and HFD lowered bone fracture toughness, while only aging reduced bone strength. At the tissue-scale, aging increased bone mineral maturity, while HFD altered bone mineralization and maturity as well as collagen structure. Cortical bone metabolism was also dysregulated with aging and HFD. Dysregulated pathways in aging and HFD were related to cellular function and viability, or glucose regulation, respectively. Aging with HFD also impacted osteoclast and adipocyte abundance and osteocyte viability. Together, these data demonstrate that HFD may exacerbate the loss of bone matrix quality and fracture resistance in aging.

## 1.0 Introduction

Bone fragility in aging is a major public health concern since the elderly (65+ years old) are the fastest growing demographic^1,2^. Direct healthcare costs for osteoporosis-related fragility fractures are projected to reach 100 billion USD by 2040^3^. This estimate is likely conservative, as 50% of fragility fractures are not fully explained by low bone mineral density (BMD) associated with osteoporosis^4^. The reason for this gap is partially explained by the alterations of bone matrix properties in aging. Bone matrix is composed of a scaffold of collagen and non-collagenous proteins, which is mineralized with hydroxyapatite. Aging changes bone matrix structure and crosslinking, which has been associated with loss of fracture resistance^5^.

Maintaining or improving the properties of bone matrix is an emerging therapeutic target^6^. However, to effectively design matrix-related therapies, it is imperative to differentiate between the effects of age and the effects of other factors that accompany aging that can modify bone matrix and, consequently, fracture resistance. The impact of lifestyle factors such as high-fat diet (HFD) is particularly important because of the potential for effective interventions and public health programming. Western diets are commonly high in saturated fats. Elderly adults are prone to eating these energy-dense, nutritionally poor diets as they are strongly associated with food insecurity, which 12% of the elderly population faces due to financial and accessibility limitations^7,8^. Aging and HFD each have detrimental effects on bone health^5,9^, but it is unknown whether HFD exacerbates the loss of bone fracture resistance and the decline in bone matrix properties in aging.

The impact of HFD on skeletal health is uncertain, partly because it is unclear whether these impacts are due to the direct effects of the diet (i.e., dietary lipids on bone health), or the excess adiposity it induces. HFD has been shown to negatively impact bone fracture resistance in rodents, including toughness^9,10^ and strength^6,9,10^. However, these prior studies used either very high fat content (≥ 60% fat)^9,10^ or long study durations with moderate (45%) fat content^6,11^. In both cases, severe metabolic disruption and excess adiposity was produced, which challenges the interpretation of the causes for decreased bone fracture resistance. Some evidence suggests that shorter-term, moderate-fat diets (≤12 weeks) in mice are sufficient to produce changes in glucose regulation and a decrease in bone strength, while avoiding large adipose tissue gains^12^. However, whether short-term, moderate-fat diet decreases bone fracture toughness and underlying matrix properties is undefined. Furthermore, whether HFD exacerbates the deleterious impacts of aging on skeletal health has not been tested.

Since the quality of bone matrix depends on the activities of bone cells, alterations in bone tissue energy metabolism are important for identifying key mechanisms underlying how aging and HFD each decrease bone fracture resistance. Both aging and HFD have been shown to alter cellular energy metabolism in non-osseus tissues, including adipose and liver tissue^13,14^. Age- and HFD-dysregulated pathways in these tissues have shown similarities, as well as sex differences, related to inflammatory response, cellular senescence, and matrix remodeling^13,14^. Bone is highly cellularized and over 90% of bone cells are osteocytes embedded within bone matrix tissue^15^. Despite *in vitro* data demonstrating that certain fats impair the activities of osteoblasts^16^ and osteocytes^17^, it is unknown whether bone tissue metabolism is dysregulated by HFD and whether these impacts would be shared or distinct from those of aging. These data are useful for identifying metabolic factors that participate in the development of bone fragility, whether produced by aging, HFD, or both.

The purpose of this study was to test the hypothesis that short-term, moderate HFD would worsen the loss of bone fracture resistance with aging in female and male C57BL/6JN mice, but that aging and HFD would have distinct effects on bone matrix and cortical bone metabolism. We found that HFD and aging each negatively affected bone fracture toughness, such that aged mice on HFD had the least tough bone. By contrast, bone strength decreased with aging but not with HFD. At the tissue-scale, the impacts of HFD and aging on bone matrix had important differences. Cortical bone tissue metabolism was dysregulated by both HFD and aging, revealing that each factor differently impacts pathways relevant to cellular energy metabolism and health. Together, these results demonstrate that short-term HFD negatively impacts skeletal health and may exacerbate the loss of bone fracture resistance in aging.

## 2.0 Results

### 2.1 HFD altered glucose metabolism but only males gained body mass

Body mass change in response to HFD varied by sex, age, and diet (sex*age*diet interaction, p < 0.001). Young (5-months-old) males gained the most body mass (Figure 1a, Supplementary Figure S1). Epididymal adipose depot mass also showed interactive effects between sex and diet (p < 0.001) and age and diet (p = 0.037), with the largest increase in HFD males compared to low-fat diet (LFD) males (+58.2%, adjusted p < 0.001, Figure 1b). Only age affected adiposity in females with aged females having higher adipose depot mass compared to young females (+56.3%, adjusted p < 0.001). Young HFD mice had heavier adipose depots than young LFD mice (+137%, adjusted p = 0.001), while aged (22-month-old) showed no significant difference in adipose depot mass between HFD and LFD.

**Figure 1.**
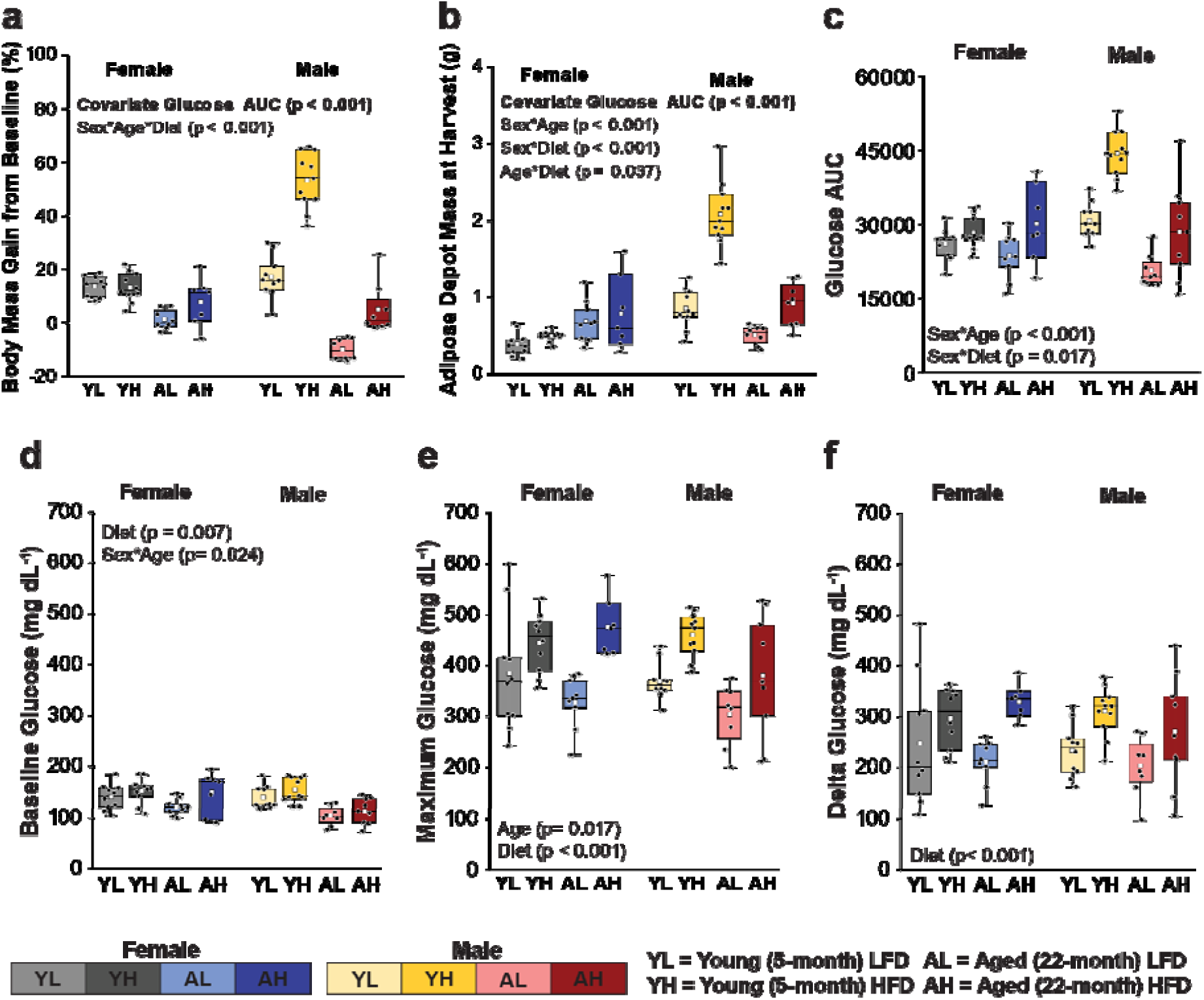
Eight weeks of high-fat diet disrupted glucose metabolism for young and aged mice of both sexes but only increased mass for males. (a) Percent mass change from the start to end of the diet intervention and (b) the reproductive adipose depot mass at harvest. (c) The area under the intraperitoneal glucose tolerance testing (ipGTT) curves was termed as the glucose area under the curve (AUC), (d) the first time point (0 min.) following fasting was the baseline glucose level and (e) the maximum glucose was the maximum glucose reading during the ipGTT. (f) Delta glucose was calculated as the difference between the maximum and baseline points. Boxplots illustrate the interquartile range (box), the minimum and maximum values (whiskers), the median value (bar across the box), the mean value (white square), and individual data points (black circles). Significant p-values from 3-way ANOVA and pairwise post-hoc tests are noted on the plots. For all models unrelated to ipGTT, the glucose AUC was considered as a covariate. If the covariate was not significant, the model was rerun without it.

Measures of glucose metabolism were significantly affected by HFD (p < 0.05 for all, Figure 1c-f, Supplementary Figure S2). Intraperitoneal glucose tolerance testing (ipGTT) glucose AUC exhibited a significant interaction between sex and diet (p = 0.017), with HFD males having greater glucose AUC than LFD males (+41.6%, adjusted p < 0.001, Figure 1c). Females showed no difference in glucose AUC between diets. Baseline, maximum, and delta (maximum – baseline) glucose levels were higher for HFD compared with LFD as a main effect (Baseline: +11.4%, p = 0.007; Maximum: +27.2%, p < 0.001; Delta: +35.5, p < 0.001, Figure 1d-f). As glucose metabolism is known to have an influence on skeletal metabolism^18^, glucose area under the curve (AUC) was used as a covariate in statistical models and removed if not significant. Notably, glucose AUC had strong positive correlations with mass gain (Pearson r = 0.85) and adiposity (Pearson r = 0.81). Full mass and ipGTT results can be found in Supplementary Table S1. These data show that the glucoregulatory response to HFD was sexually dimorphic, which also provides an opportunity to begin separating the effects of HFD from adiposity.

### 2.2 Systemic response to HFD-induced stress depends on sex and age

Serum biomarkers were assessed to determine the independent and interactive effects of sex, diet, and age on systemic metabolism and inflammatory homeostasis. The circulating factors assessed in this study regulate bone turnover, bone cell differentiation, glucoregulation, and feeding behavior^19,20^. The adipokine leptin exhibited a three-way interaction (p = 0.018), where young HFD male mice had over 20-fold higher levels compared to all other groups (Figure 2a). Three-way interactions were also found for the adipokine resistin (p = 0.011, Figure 2b) and the angiogenic factor vascular endothelial growth factor (VEGF, p = 0.032). Aged HFD males had lower resistin levels compared to aged LFD males (-37.0%, p = 0.001), with no effect of age or diet on resistin in females. Diet did not impact VEGF in young males and females, or in aged females; however, aged LFD males had higher VEGF levels than aged HFD males (+34.0%, p = 0.002). Aged females, independent of diet, had higher VEGF levels compared to young females (LFD: +35.9%, HFD: +51.7%, p < 0.002 for both), while aged LFD males had higher VEGF compared to young LFD males (+43.3%, p = 0.005, Figure 2c). Leptin, resistin, and VEGF are expected to change in response to altered glucoregulation and these results demonstrate that the response of these cytokines to diet depends on age and sex.

**Figure 2.**
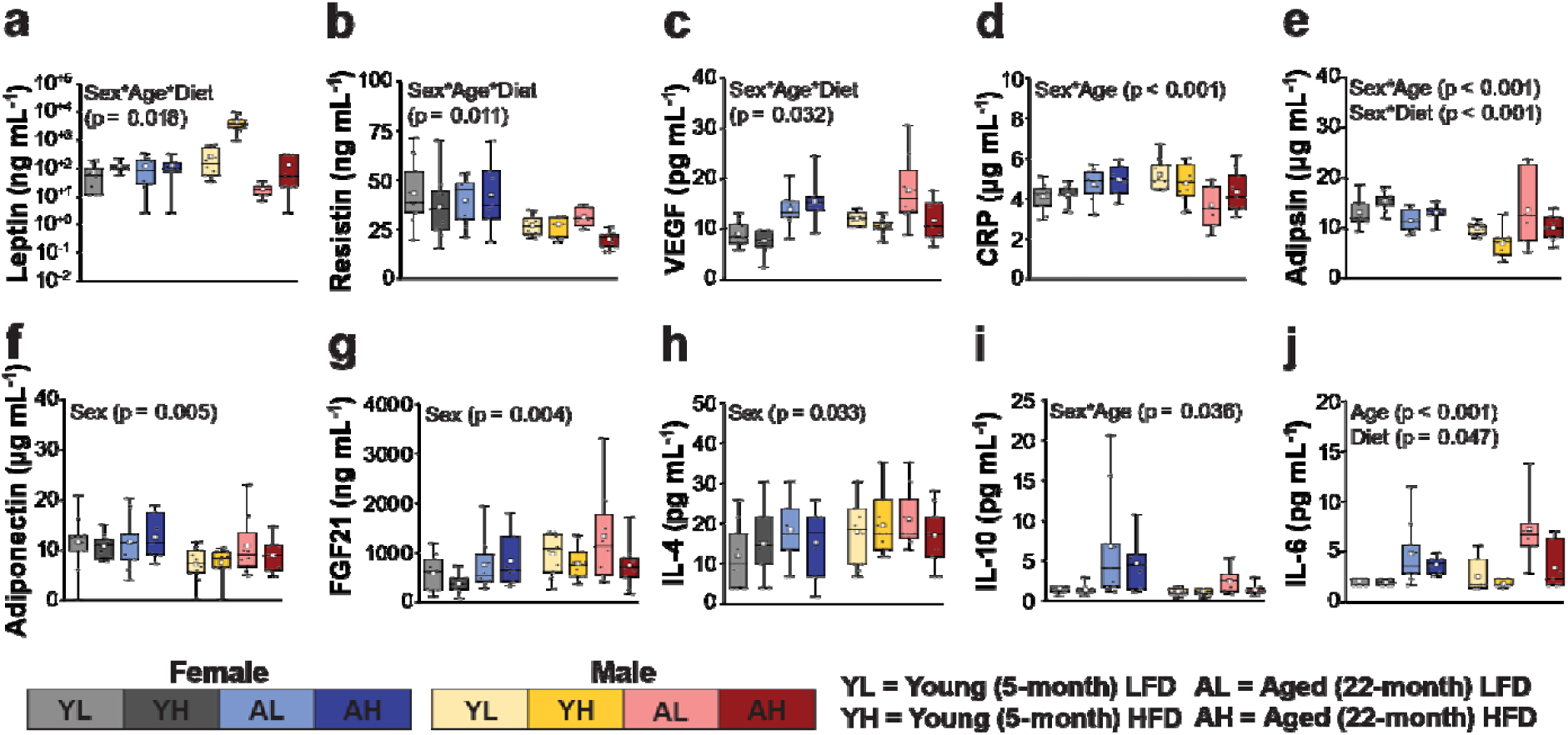
The systemic inflammatory response to high-fat diet depends on sex and age. Levels of inflammatory biomarkers: (a) leptin, (b) resistin, (c) vascular endothelial growth factor (VEGF), (d) C-reactive protein (CRP) (e) adipsin, (f) adiponectin, (g) fibroblast growth factor 21 (FGF21), (h) interleukin 4 (IL-4), (i) IL-10, and (j) IL-6. Boxplots illustrate the interquartile range (box), the minimum and maximum values (whiskers), the median value (bar across the box), the mean value (white square), and individual data points (black circles). Significant p-values from 3-way ANOVA and pairwise post-hoc tests are noted on the plots. For all models, the glucose area under the curve (AUC) was considered as a covariate. If the covariate was not significant, the model was rerun without it.

C-reactive protein (CRP) showed an interaction between sex and age (p < 0.001) but no diet effect. Young males had higher CRP levels than aged males (+19.5%, p = 0.001), but no age effect was observed in females (p = 0.017). Young females had lower CRP levels compared to young males (-16.7%, p = 0.002), while aged females had higher levels compared to aged males (+17.0%, p = 0.007) (Figure 2d). These results show that CRP, a cytokine that would be expected to increase with adiposity^21^, does not change with diet but does depend on age in a manner that is sex-dependent.

HFD affected circulating levels of the adipokine adipsin, with significant interactions between HFD and sex (p = 0.001) and age and sex (p = 0.001). HFD males had lower adipsin levels compared to LFD males (-28.1%, p = 0.001), and HFD females had higher levels compared to HFD males (+42.3%, p < 0.001). Young females had higher adipsin levels compared to young males (+41.8%, p < 0.001, Figure 2e). There were notable sex differences in several markers. Adiponectin was higher in females (+26.0%, p = 0.005, Figure 2f), while Fibroblast growth factor 21 (FGF21) and interleukin-4 (IL-4) were elevated in male mice (FGF21: +35.7%, p = 0.004; IL-4: +31.6%, p = 0.033, Figure 2g and h). IL-10 demonstrated a sex and age interaction (p = 0.036). Aged females had elevated IL-10 levels compared to aged males (+58.9%, p < 0.001) and young females (+196%, p < 0.001), with no significant difference between age groups in males (Figure 2i). IL-6 was higher in aged compared to young mice, as expected^22^ (+83.9%, p < 0.001) and, unexpectedly, lower in HFD compared to LFD groups (-16.7%, p = 0.047)^23,24^ (Figure 2j). Full results of serum biomarker assays can be found in Supplementary Table S2. In summary, HFD influenced some cytokines that have an expected association with altered glucoregulation, including adipsin^25^ and IL-6^23^. However, the relationship between these cytokines and diet did not reflect all the outcomes seen in obesity.

Specifically, IL-6 decreased with HFD and adipsin only increased for males. Furthermore, there were several cytokines associated with high adiposity that did not change with diet (IL-10, IL-4, and FGF21)^26–28^.

### 2.3 HFD has minimal impact on measures of cortical and trabecular microarchitecture

Cortical bone expanded in an interactive manner between age and sex (p = 0.023), as evidenced by the marrow area expanding in both sexes (aged vs. young females: +38.2%; aged vs. young males: +40.2%, adjusted p < 0.001 for both). Cortical bone also thinned with age (cortical thickness: -12.0%, p < 0.001), as expected^29^. HFD did not influence cortical bone size (e.g. minimum moment of inertia, p > 0.05, Figure 3a-e) or cortical thickness (p = 0.072, Figure 3b). Cortical area was reduced by HFD (-3.5%, p = 0.033) when including a significant outlier, but became insignificant when excluding this outlier (p = 0.052). Cortical porosity was unaffected by diet but demonstrated an interaction between sex and age (p = 0.035), with males having more porous bone than females at both ages (young: +42.2%, aged: +22.7%, adjusted p < 0.001 for both, Figure 3c).

**Figure 3.**
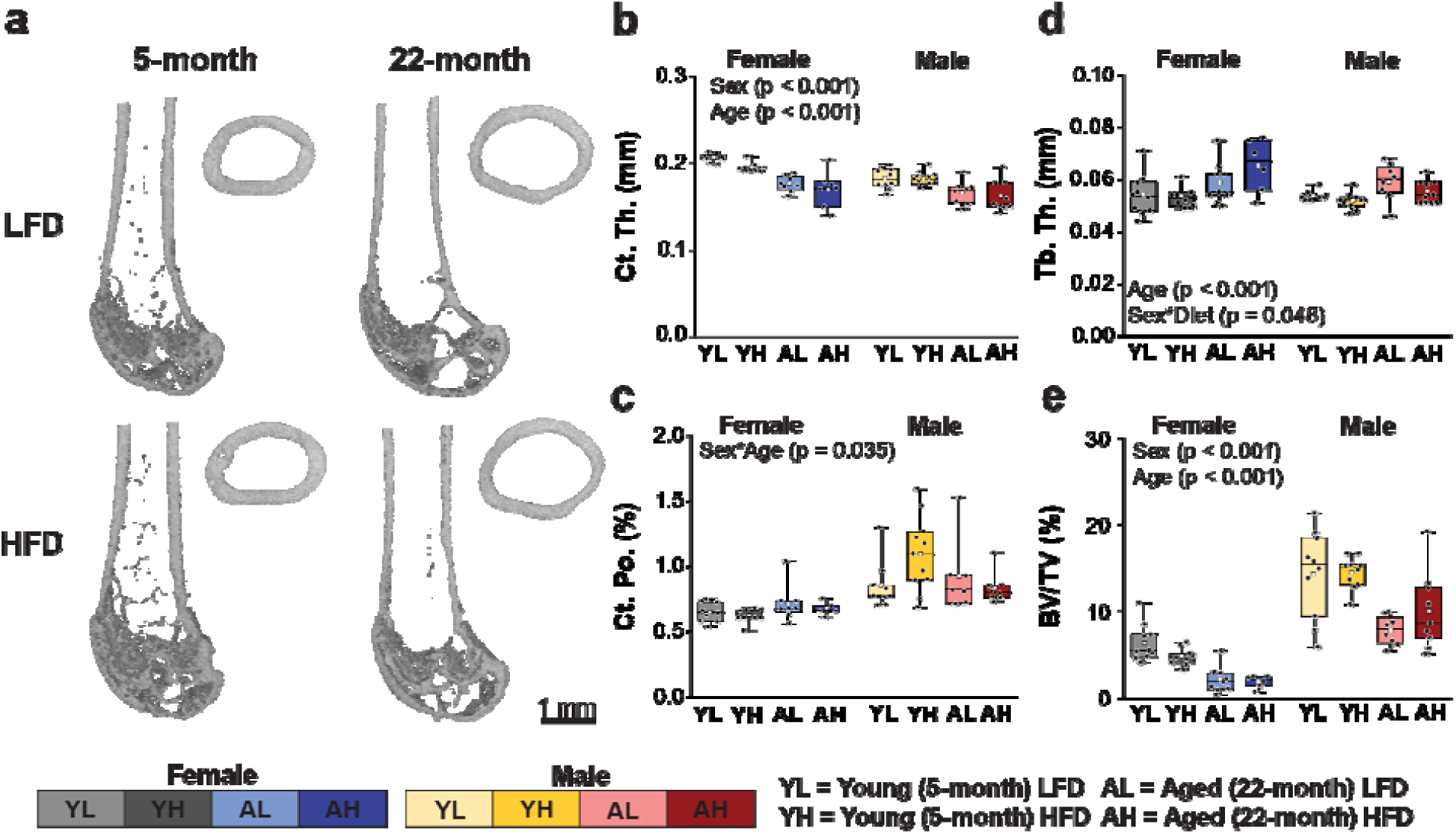
High-fat diet did not affect most measures of cortical and trabecular microarchitecture. (a) Representative micro-tomographic images for female groups. Cortical bone measures demonstrated by (b) cortical thickness (Ct. Th.) and (c) cortical porosity (Ct. Po.). Trabecular bone measures demonstrated by (d) trabecular thickness (Tb. Th.) and (e) trabecular bone volume (BV/TV). Boxplots illustrate the interquartile range (box), the minimum and maximum values (whiskers), the median value (bar across the box), the mean value (white square), and individual data points (black circles). Significant p-values from 3-way ANOVA and pairwise post-hoc tests are noted on the plots. For all models, the glucose area under the curve (AUC) was considered as a covariate. If the covariate was not significant, the model was rerun without it.

Few trabecular microarchitecture measures were affected by diet. BMD had a significant interaction between sex and diet (p = 0.010) with HFD females having a lower BMD compared to LFD females (-15.5%, adjusted p = 0.014). There was no difference of BMD between diets in males. Trabecular thickness was affected by the interaction of sex and HFD (p = 0.048), but HFD did not increase trabecular thickness within each sex. HFD females had thicker trabeculae compared to HFD males (+9.7%, adjusted p = 0.008, Figure 3d). No other measures of trabecular microarchitecture were impacted by HFD. However, trabecular bone volume, trabecular number, and connectivity density were lower in aged and female mice compared to young and male mice, respectively. Complete data for femur cortical geometry and trabecular microarchitecture are reported in Supplementary Table S3. This data demonstrates that short-term HFD has little effect on microarchitecture regardless of the degree of adiposity developed.

### 2.4 HFD decreases fracture toughness but not bone strength

Bone fracture resistance depends on fracture toughness and strength^30^. HFD mice had lower bone tissue intrinsic fracture toughness compared to LFD mice (Kc_init_, -15.1%, p = 0.023, Figure 4a). There was also a trend (p = 0.056) for lower fracture toughness at maximum loading (Kc_max_) for HFD compared to LFD groups. HFD did not interact with age or sex effects on fracture toughness. However, Kc_max_ demonstrated an interaction between sex and age (p = 0.019), where aged females had lower fracture toughness than young females (-23.2%, adjusted p = 0.002, Figure 4b). Males did not have significant differences between age groups for Kc_max_. Kc_init_ was higher for young mice (+15.9%, p = 0.017) but showed no sex effect.

**Figure 4.**
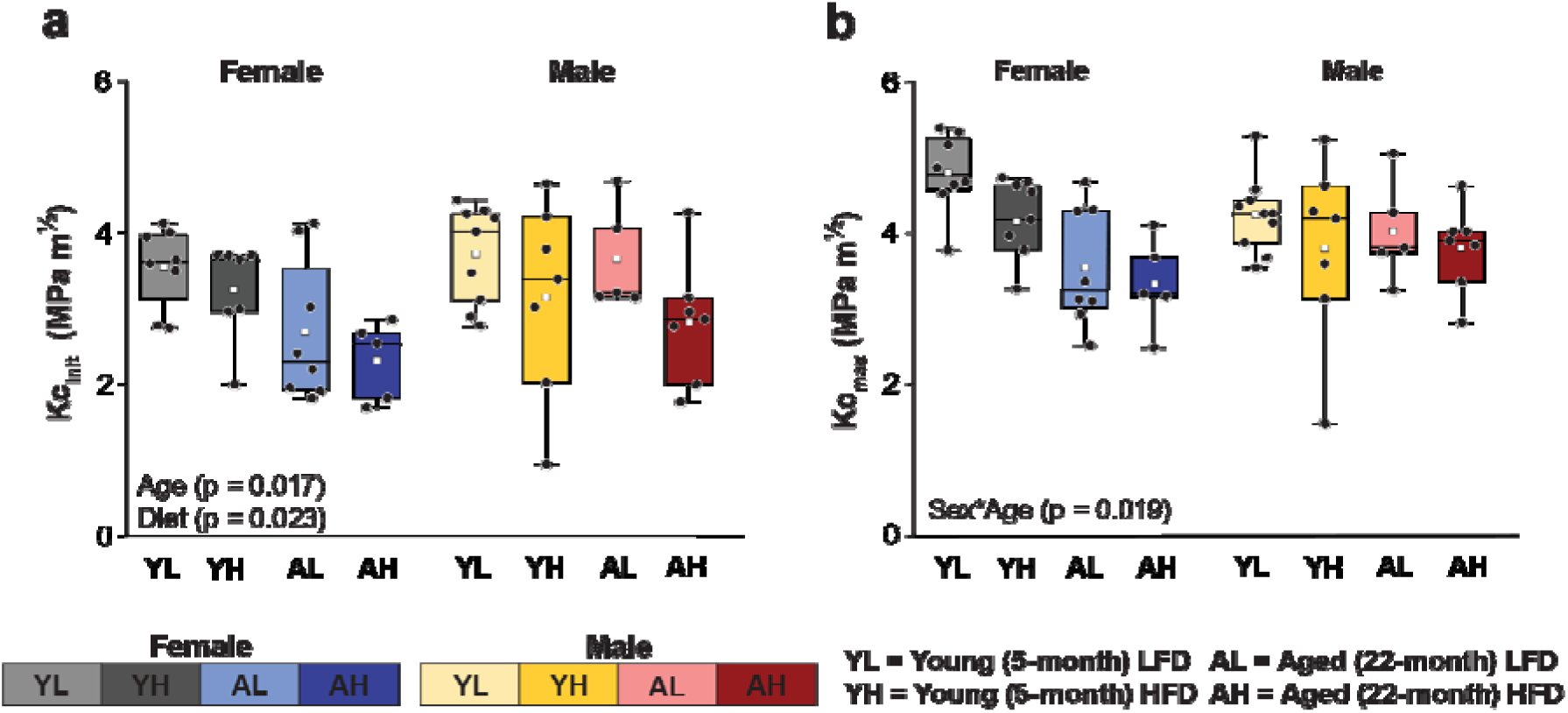
Aging and high-fat diet reduce bone intrinsic toughness to fracture. (a) Intrinsic fracture toughness (Kc_init_) and (b) fracture toughness at maximum load (Kc_max_). Boxplots illustrate the interquartile range (box), the minimum and maximum values (whiskers), the median value (bar across the box), the mean value (white square), and individual data points (black circles). Significant p-values from 3-way ANOVA and pairwise post-hoc tests are noted on the plots. For all models, the glucose area under the curve (AUC) was considered as a covariate. If the covariate was not significant, the model was rerun without it.

Bone strength showed a significant interaction between sex and diet (p = 0.021), with HFD females having stronger bones than HFD males (+16.9%, adjusted p < 0.001). LFD females and males did not differ, and diet groups within each sex were not significantly different. As expected, strength was significantly lower in aged mice (-30.0%, p < 0.001)^31,32^. HFD did not impact the elastic modulus, but elastic modulus was affected by both age and sex, where females had higher moduli than males (+23.7%, p < 0.001) and aged mice had lower moduli than young mice (-34.0%, p < 0.001). Yield strength was not affected by diet or sex but was decreased in aged mice compared to young mice (-44.4%, p < 0.001). Complete measurements from three-point bending and notched fracture toughness testing are reported in Supplementary Table S4. These data show that HFD and age have different effects on bone fracture resistance. Specifically, HFD lowers fracture toughness, while aging lowers both fracture toughness and strength.

### 2.5 HFD alters local, but not global, measures of bone turnover

The percent of mineralizing matrix on the endocortical surface (E. MS/BS, Figure 5a) demonstrated main effects of diet (p = 0.019) and sex (p < 0.001). HFD mice had less mineralizing endocortical surface than LFD mice (-33.4%), and males had less mineralizing endocortical surface than females (-69.4%). In contrast, the periosteal mineralizing surface (P. MS/BS, **F**igure 5b) showed only an age effect (p < 0.001) where aged mice had less mineralizing surface than young mice (-70.8%). However, age and sex may have an interactive effect on the periosteal mineralizing surface, but it did not reach statistical significance (p = 0.053).

**Figure 5.**
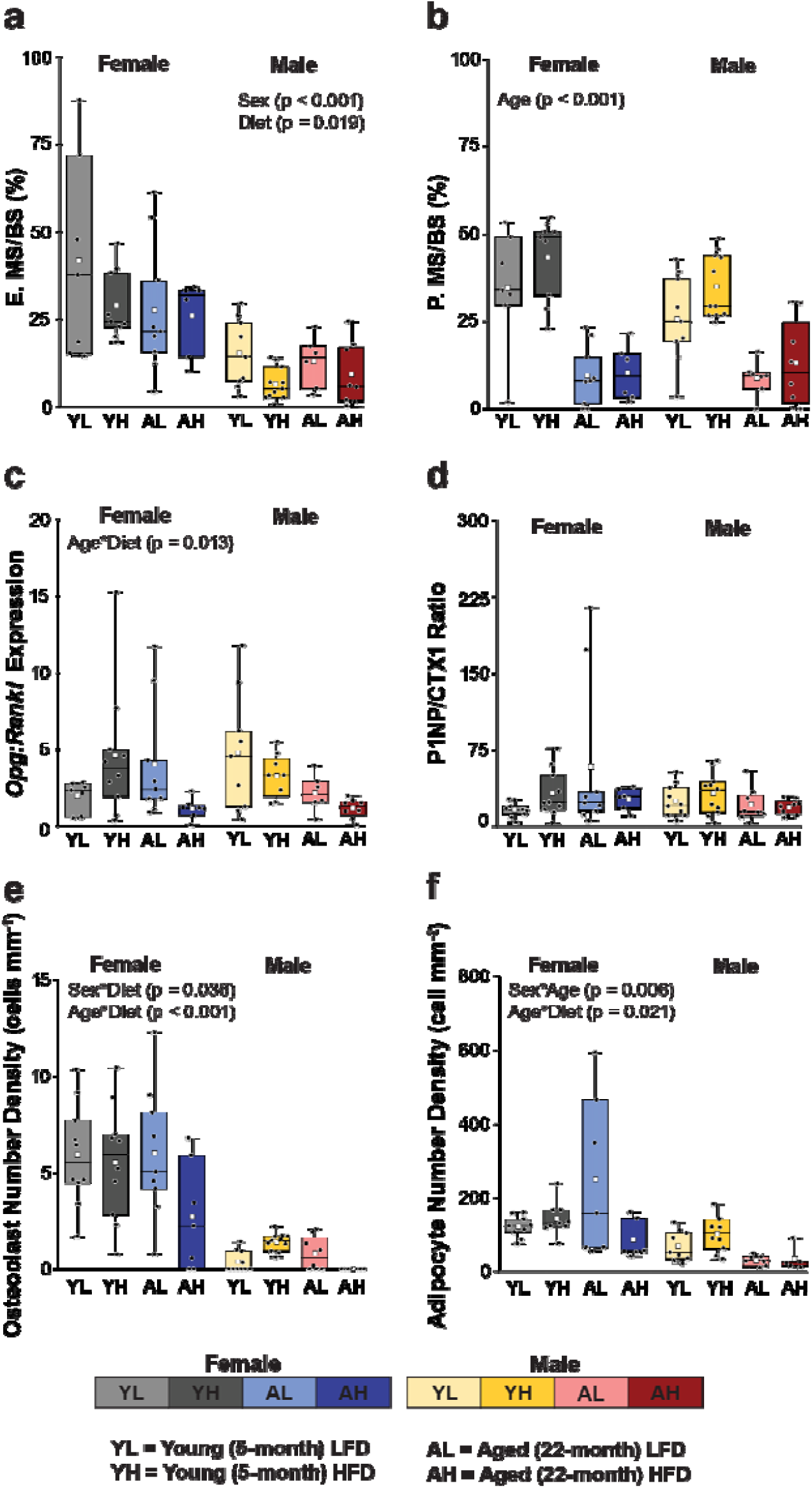
High-fat diet and aging alter bone turnover. (a) Endocortical mineralizing surface (E. MS/BS) and (b) periosteal mineralizing surface (P. MS/BS) evaluated from quantitative histomorphometry are local measures of bone mineralization. (c) The whole-bone *Opg:Rankl* ratio gene expression from marrow-flushed tibiae and (d) blood serum P1NP:CTX1 ratio are global markers of bone turnover. (e) Tartrate resistant acid phosphatase (TRAP)-positive osteoclast number density and (f) bone marrow adipose tissue (bMAT) adipocyte number density from the tibia metaphysis are measures related to cellular activity necessary for bone turnover. Boxplots illustrate the interquartile range (box), the minimum and maximum values (whiskers), the median value (bar across the box), the mean value (white square), and individual data points (black circles). Significant p-values from 3-way ANOVA and pairwise post-hoc tests are noted on the plots. For all models, the glucose area under the curve (AUC) was considered as a covariate. If the covariate was not significant, the model was rerun without it.

Gene expression of *Opg:Rankl* from flushed tibia had an interactive effect between age and diet (p = 0.013), with aged HFD mice having lower expression compared to aged LFD mice (-63.2%, adjusted p = 0.009) and young HFD mice (-72.4%, adjusted p < 0.001, Figure 5c). The serum ratio of P1NP:CTX1, a global biomarker of bone formation and resorption was unaffected by diet, age, or sex (Figure 5d).

HFD also affected bone resorption. Osteoclast number density, indicated by tartrate resistant acid phosphatase (TRAP)-positive multinucleated cells, showed an interaction between diet and age (p < 0.001), where aged HFD mice had lower osteoclast number density compared to aged LFD mice (-81.0%, p < 0.001, Figure 5e). Diet had no impact on osteoclast density in young mice. Osteoblasts and adipocytes differentiate from mesenchymal stem cells and both aging and HFD promote adipogenic differentiation^33,34^. There was no diet main effect on bone marrow adiposity, but an interaction between age and diet (p = 0.021) demonstrated that aged mice with HFD had lower adipocyte density than young HFD mice (-61.7%, adjusted p < 0.001, Figure 5f). Complete measurements from quantitative histomorphometry, serum biomarker, qRT-PCR, and histology are reported in Supplementary Tables S5-S7.

These data demonstrate that the effect of HFD on bone turnover is detectable at the local, but not global, levels. Furthermore, this local effect is site-specific, and is more pronounced at the endocortical as compared with periosteal surfaces. Whereas HFD had an effect on bone formation at all ages, there was only an impact of diet on osteoclast and adipocyte counts in aged mice.

### 2.6 HFD reduces osteocyte viability and affects some measures of lacunar-canicular system turnover gene expression

Since aging decreases osteocyte viability^35–38^, terminal deoxynucleotidyl transferase dUTP nick end labelling (TUNEL) staining was used to quantify osteocytes exhibiting DNA fragmentation. The percentage of TUNEL-positive osteocytes showed interaction between age and diet (p = 0.024), where aged HFD mice had more TUNEL-positive osteocytes compared to aged LFD (+75.3%, adjusted p = 0.003) and young HFD mice (+62.5%, adjusted p = 0.005, Figure 6a). Osteocyte number density showed an interaction between sex and diet (p = 0.014), where HFD males had more osteocytes than LFD males (+45.1%, adjusted p = 0.001) but females did not differ between HFD and LFD groups (Figure 6b). There was also an interaction between sex and age (p = 0.001), where aging reduced osteocyte number density but only for males (-36.4%, adjusted p < 0.001). Osteocyte histology results are reported in Supplementary Table S8.

**Figure 6.**
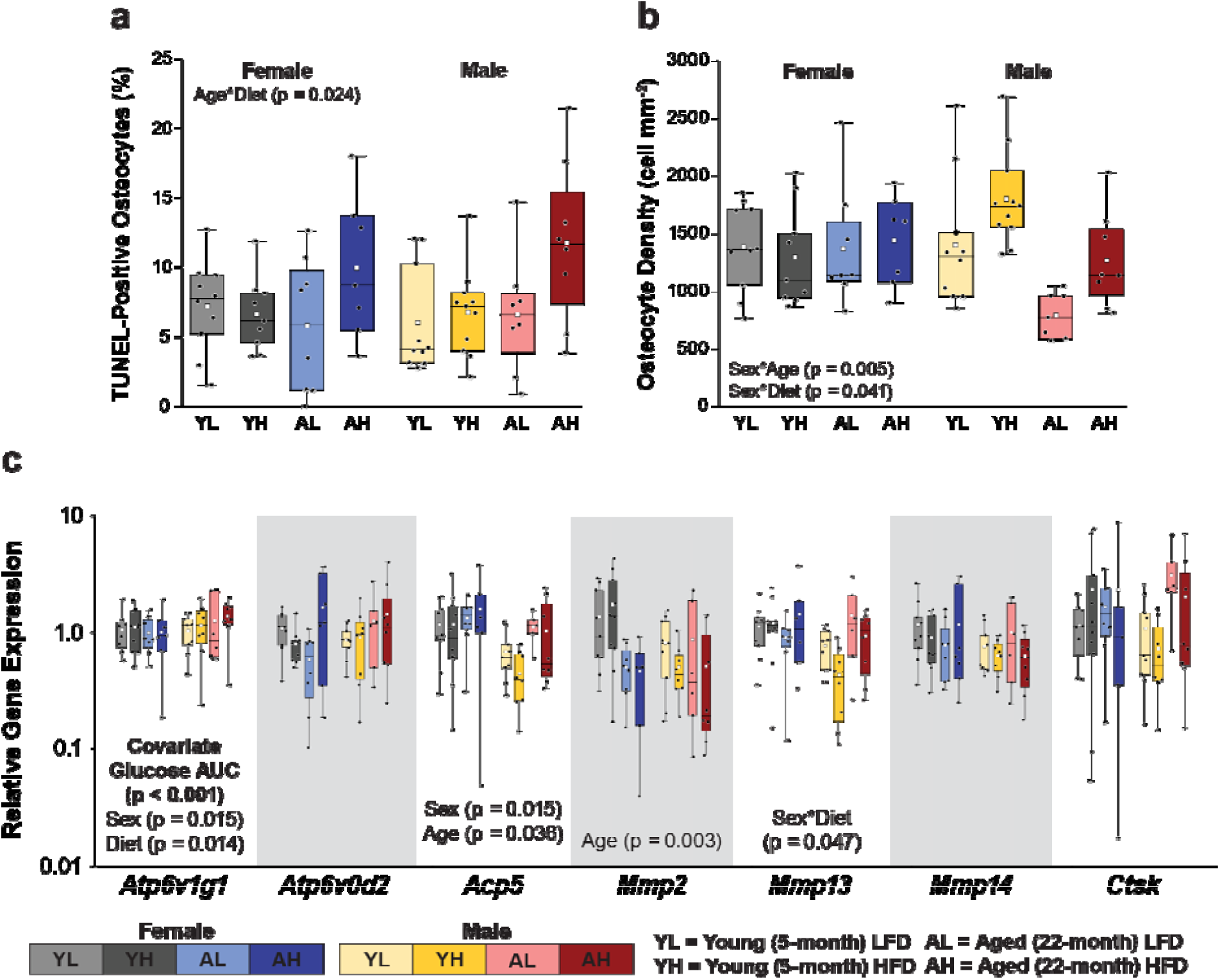
High-fat diet and aging alter osteocyte viability and gene expression related to lacunar-canicular system (LCS) turnover. (a) Osteocytes positive for terminal deoxynucleotidyl transferase dUTP nick end labelling (TUNEL), a marker of apoptosis. (b) Osteocyte number density. (c) Relevant genes expressed during LCS bone resorption measured in marrow-flushed tibiae. Boxplots illustrate the interquartile range (box), the minimum and maximum values (whiskers), the median value (bar across the box), the mean value (white square), and individual data points (black circles). Significant p-values from 3-way ANOVA and pairwise post-hoc tests are noted on the plots. For all models, the glucose area under the curve (AUC) was considered as a covariate. If the covariate was not significant, the model was rerun without it.

In flushed tibiae, expression of genes related to osteocyte lacunar-canalicular (LCS) resorption showed age effects, where *Mmp2* was lower in aged mice (-52.1%, p = 0.003), while *Acp5* was higher in aged mice compared to young mice (+47.3%, p = 0.036). Gene expression was also affected by diet. HFD mice, covarying for glucose AUC (p < 0.001), had higher expression of vacuolar ATPase H+ transporting *v1* subunit *g1* compared with LFD (*Atp6v1g1*, +37.6%, p = 0.014) and had higher expression in males compared to females (+32.2%, p = 0.015). *Mmp13* showed a significant interaction between sex and diet (p = 0.047), with HFD females having higher *Mmp13* expression compared to HFD males (+48.5%, adjusted p = 0.008). HFD did not impact *Mmp13* gene expression compared to LFD for either sex (Figure 6c). *Mmp14*, *Mmp2*, *Acp5*, *Atp6v0d2,* and *Ctsk* were not significantly impacted by diet. Full LCS turnover gene expression results are reported in Supplementary Table S9. These data show that HFD reduces osteocyte viability in aged but not young mice. However, the impact of HFD on LCS turnover was mixed. Some markers were higher in HFD groups (*Atp6v1g1*), others depended on an interaction between sex and HFD (*Mmp13*), while others did not depend on diet but were influenced by age or sex (e.g. *Acp5*).

### 2.7 High-fat diet alterations to bone matrix at the endocortical surface are associated with decreased fracture toughness

Fluorescent AGEs (fAGEs) are matrix molecules that promote adducts and non-specific crosslinks between matrix fibrils, disrupting intrinsic toughening mechanisms in bone^39^. There was a three-way interaction between sex, age, and diet (p = 0.033). However, post-hoc correction for multiple comparisons demonstrated that fAGEs were not significantly different between groups (all comparisons had greater p than new significance level p = 0.006, Supplementary Table S10), nor were they strongly correlated with the decrease of Kc_init_ in aged or HFD at either bone surface (Figure 8a and b).

Bone matrix properties were evaluated at the periosteal surface and in polished cross-sections using Raman spectroscopy under hydrated condition to mimic *in vivo* properties. At the periosteal surface, sex and age differences were evident in mineral quantity and maturity. Males had lower median mineral:matrix than females (-12.1%, p < 0.001), and aged mice had higher mineral results of maturity, as estimated by the median carbonate:phosphate ratio, than young mice (+11.6%, p < 0.001). Bone median mineral crystallinity had an interaction between sex and age p = 0.016), where aged males, but not females, had modestly higher median mineral crystallinity compared to young counterparts (+2.5%, adjusted p = 0.018). There was no impact of diet on the median of either periosteal matrix measures (Figure 7 Row 1).

**Figure 7.**
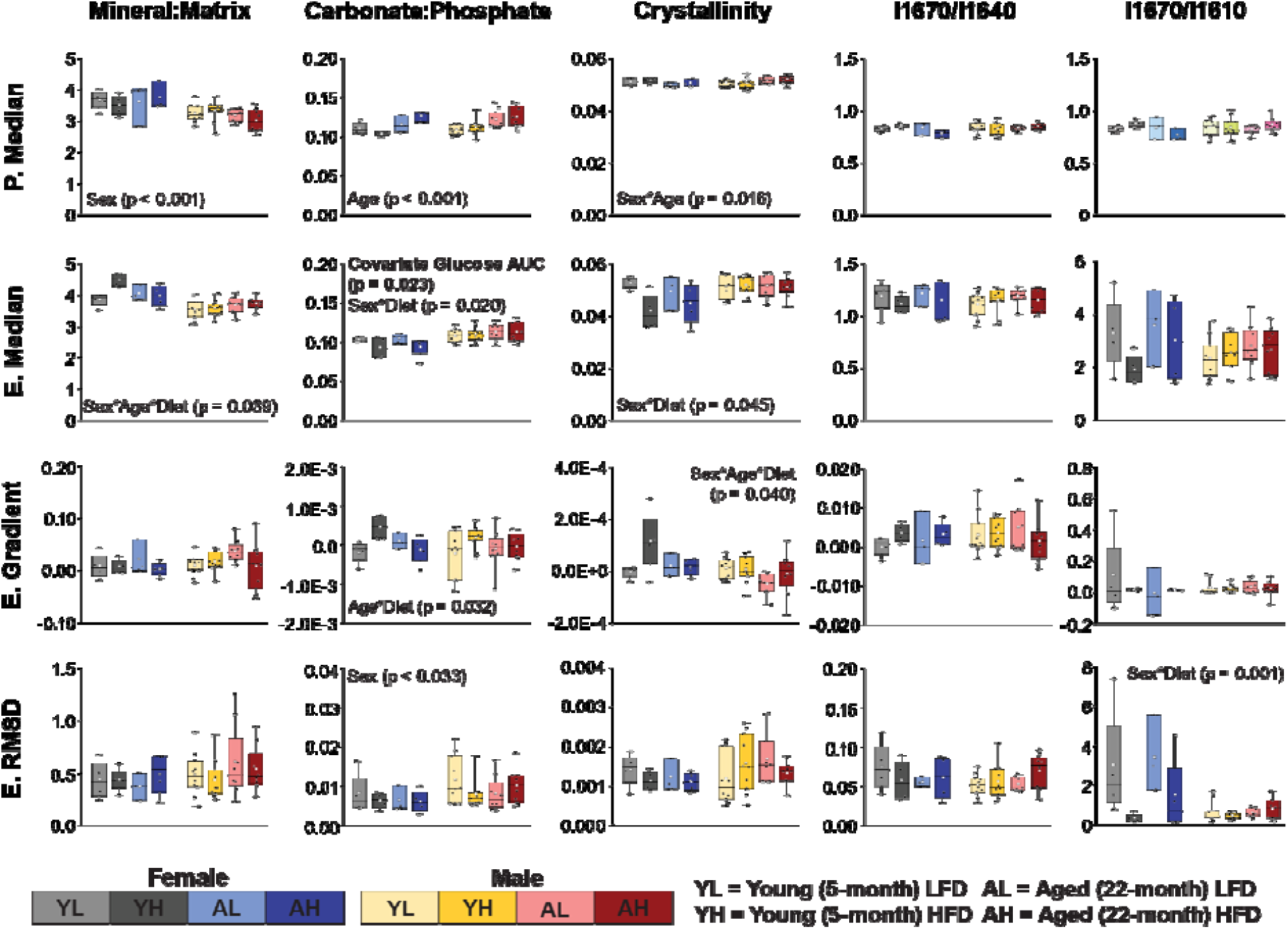
Bone matrix composition depends on high-fat diet in an age- and sex-dependent manner. The measurements of matrix maturity from hydrated Raman spectroscopy for the periosteal surface and the endocortical region (anterior-lateral location). Row 1 – the median measures for the periosteal surface (P. Median). Row 2 – the median measures for the endocortical region (E. Median). Row 3 – the gradient (slope of linear fit) of the measures across the endocortical region (E. Gradient). Row 4 – Estimate of the root mean square deviation (RMSD) from linear fit residuals of each measure across the endocortical region (E. RMSD). Boxplots illustrate the interquartile range (box), the minimum and maximum values (whiskers), the median value (bar across the box), the mean value (white square), and individual data points (black circles). Significant p-values from 3-way ANOVA and pairwise post-hoc tests are noted on the plots. For all models, the glucose AUC was considered as a covariate. If not significant, the model was rerun without the covariate.

**Figure 8.**
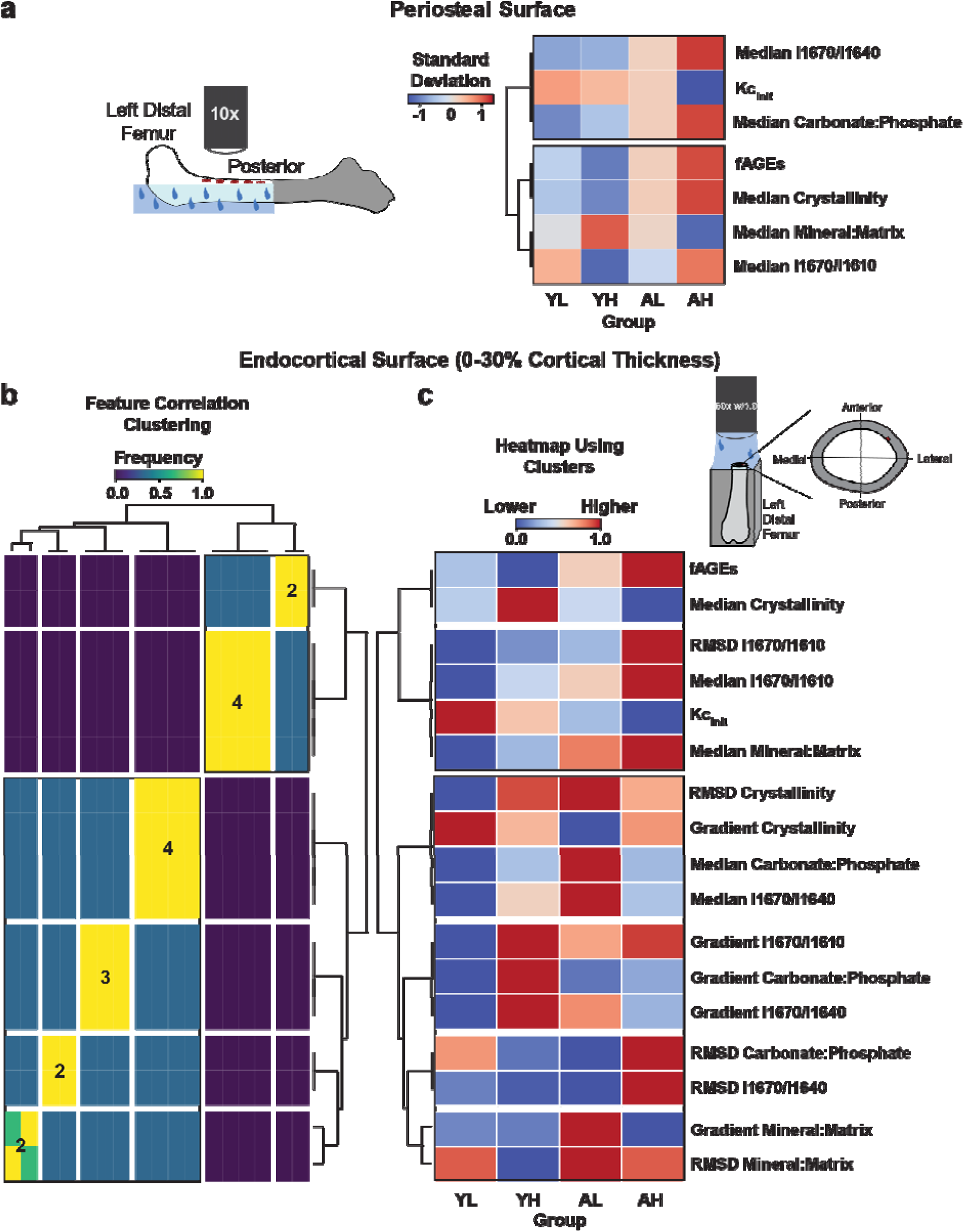
Bone matrix alterations at the endocortical surface are associated with decreased fracture toughness. (a) Heatmap of periosteal hydrated Raman spectroscopy collected with the set up illustrated on the left distal femur. Measures for male samples were clustered by magnitude of absolute correlation. (b) A frequency map of ensemble clustering with cluster optimization (ECCO) clustering solutions of absolute correlations from endocortical Raman spectroscopy measures in males. c) ECCO clusters were then used to create a heatmap of matrix measures. YL = Young (5-month) low-fat diet, YH = Young (5-month) high-fat diet, AL = Aged (22-month) low-fat diet, AH = Aged (22-month) high-fat diet, fAGEs = fluorescent advanced glycation end products, Kc_init_ = Intrinsic fracture toughness, Gradient = Slope of the linear fit of the data, RMSD = Root mean square deviation of the data about the linear fit.

Quantitative histomorphometry (Figure 5a) indicated HFD altered bone mineralization at the endocortical surface, which prompted us to examine how the spatial variance of bone matrix properties may be altered within this region of interest. One aspect of spatial variance is the slope of change for a measure with respect to radial distance from the endocortical surface (i.e., gradient). Another aspect of spatial variance is the variability of the data (i.e., root mean square deviation, RMSD; Supplementary Figure S3). Raman spectroscopy showed sex and diet effects on matrix quality in the endocortical region (i.e. first 30% of bone thickness from endocortical surface, Figure 7 Row 2-4). Interactions between sex and diet were significant for median mineral crystallinity and RMSD of collagen secondary structure (RMSD I1670/I1610) (p = 0.045; p = 0.001 respectively). HFD females had lower median mineral crystallinity compared to LFD females (-15.4%, adjusted p = 0.017) as well as lower variability of collagen secondary structure, as assessed by decreased RMSD I1670/I1610 (-85.1%, adjusted p < 0.001). Males showed no significant diet differences in either measure. Median mineral:matrix had a three-way interaction (p = 0.039). After multiple comparison corrections, young HFD females had higher median mineral:matrix content compared to their both young LFD (+16.8%, adjusted p = 0.002). There was no impact of diet on these measures in male mice. Full Raman spectroscopy results are in Supplementary Table S11.

Since the intrinsic fracture toughness of bone tissue was lower with HFD and aging, and multiple, often intercorrelated, features of matrix properties contribute to intrinsic toughening, we utilized clustering techniques based on the strength of absolute correlations to assess the association of matrix properties with fracture toughness (Kc_init_). Only male results were clustered due to the relatively limited availability of female data. At the periosteal surface, the carbonate:phosphate ratio and collagen secondary structure (I1670/I1640) measures had the strongest correlations (inverse) with fracture toughness as evident through representation in the same cluster (Figure 8a). At the endocortical surface, the median collagen secondary structure (I1670/I1610), RMSD of collagen secondary structure (RMSD I1670/I1610), and the median mineral:matrix ratio had the strongest (inverse) correlations to fracture toughness. Decreased fracture toughness was associated with higher values for these measures. fAGEs were not within the closest clusters with Kc_init_ for either the periosteal or endocortical region of interest (Figure 8b).

Together, these data show that HFD and aging differently impact bone matrix. HFD alters endocortical bone mineral and, for females, the RMSD of collagen secondary structure. By contrast, aging increases periosteal bone mineral maturity, but not mineral content or collagen structure. The variation of fracture toughness seen across males in this study was more related to variation in collagen structure and mineral:matrix ratio than other measures.

### 2.8 HFD and aging differently dysregulate cortical bone metabolism

Given the robust impact of diet and age on systemic metabolism, including the liver and adipose tissue^13,14^, we were motivated to ask whether HFD also dysregulates the metabolism of cortical bone tissue. We used untargeted liquid chromatography-mass spectrometry (LC-MS) and the unbiased clustering tool ensemble clustering with cluster optimization (ECCO)^40^ to establish the impact of HFD and aging on cortical bone metabolism from marrow-flushed humeri.

Metabolomics analyses rely on clustering data. Usually, one combination of linkage and distance functions amongst many choices is chosen that define a clustering solution. This approach can potentially reduce the reproducibility of results, since other clustering methods may yield different solutions. ECCO offers an approach that computes multiple clustering solutions from different combinations of linkage and distance functions and, in this case, identified which metabolites co-occurred in 100% of clustering solutions^40^. ECCO was used for two complementary approaches. One approach identified metabolomic signatures representative of each study factor (i.e., aging, diet, and each sex). The second approach identified bioenergetic differences between pairs of groups (e.g., young HFD females vs aged HFD females).

The first approach was to use ECCO to identify the metabolites that clustered with all groups sharing a specific study factor (i.e., aging, diet, or sex), and the pathways associated with these metabolites. We then compared the metabolic signatures for distinct and shared pathways between the factors of HFD and aging. The factor of HFD was associated with 20 pathways. The following 5 pathways were distinct to the HFD signature and not overlapping with the aged signature: branched chain amino acid biosynthesis, aromatic amino acid biosynthesis, lysine degradation, terpenoid backbone biosynthesis, and linoleic acid metabolism (Figure 9a). These pathways are associated with the development and progression of glucose and insulin intolerance^41^ (Figure 9b). The factor of aging was associated with 18 pathways. The following 3 pathways were distinct to the aged signature and not overlapping with the HFD signature: pentose phosphate pathway, mannose type O-glycan biosynthesis, and ubiquinone and other terpenoid-quinone biosynthesis (Figure 9a). These pathways are linked to overall cellular functional decline and death (Figure 9b)^42–45^.

**Figure 9.**
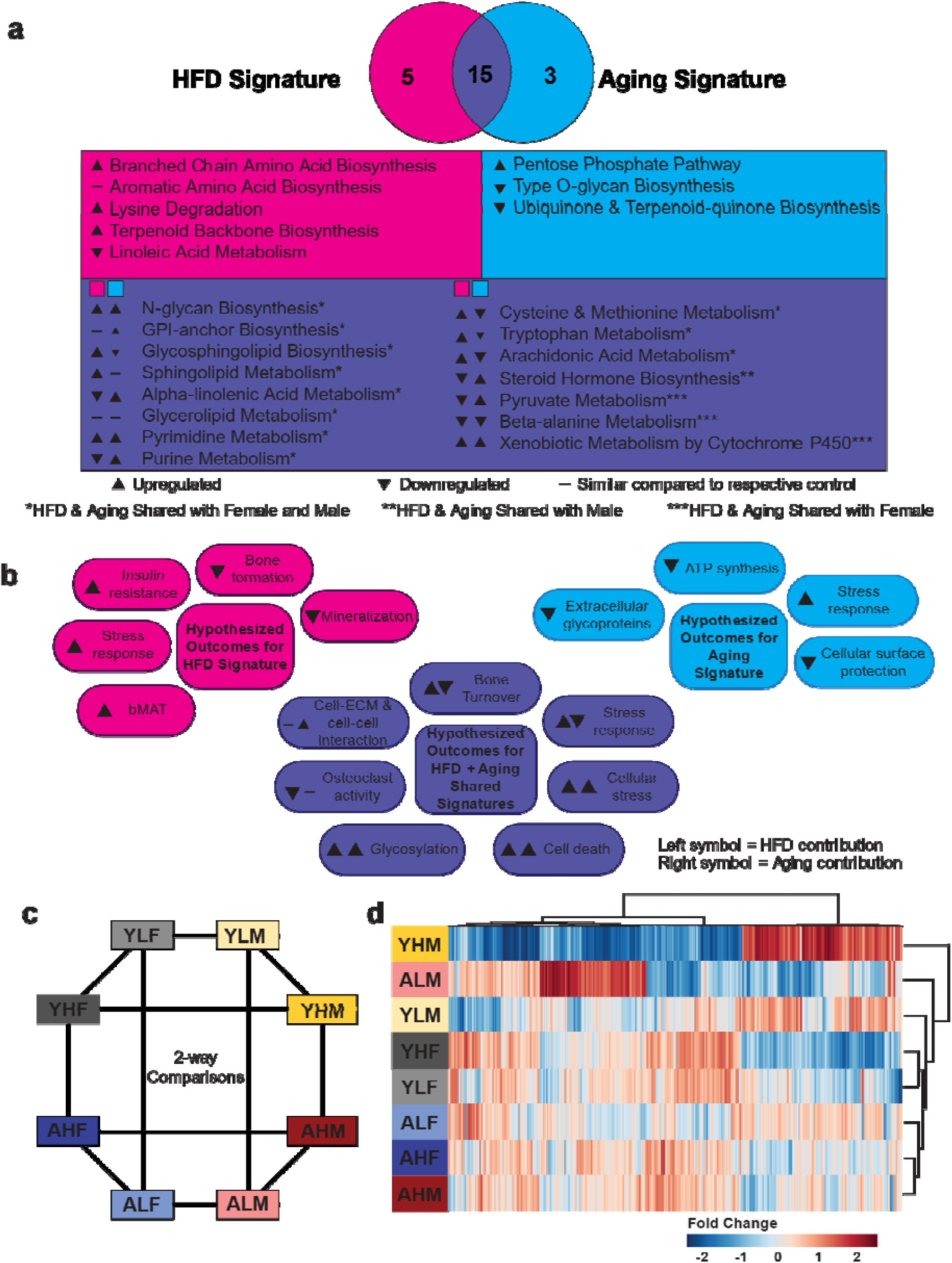
High-fat diet and aging differently dysregulate cortical bone metabolism. (a) Summary of the distinct signature pathways for high-fat diet (HFD, pink) and aged (cyan) groups and the shared overlapping signature pathways (purple) from 4-group comparisons. Signature pathways that remain shared between aged and HFD when including both sex signatures are marked by *. HFD and aged pathways shared with a particular sex signature are noted by ** = shared with males and not females or *** = shared with females and not males. Arrows indicate relative metabolite levels compared to a signature’s respective control (e.g. HFD compared to low-fat diet (LFD) levels). (b) Potential outcomes of the distinct signatures for HFD (pink), aged (cyan), and their shared overlapping pathways (purple) based on literature. (c) Diagram of all 2-group comparisons performed using ensemble clustering with cluster optimization (ECCO). (d) Heatmap of all group averages clustered by metabolite and group. YLF = Young (5-month) LFD female, YHF = Young (5-month) HFD female, AHF = Aged (22-month) HFD female, ALF = Aged (22-month) LFD female, YLM = Young (5-month) LFD male, YHM = Young (5-month) HFD male, AHM = Aged (22-month) HFD male, ALM = Aged (22-month) LFD male.

There were 15 pathways overlapping between HFD and aged signatures, with 73% conserved across sexes. Overlapping pathways included: steroid hormone biosynthesis, purine metabolism, arachidonic acid metabolism, alpha-linolenic acid metabolism, beta-alanine metabolism, cysteine and methionine metabolism, tryptophan metabolism, glycosylphosphatidylinositol (GPI)-anchor biosynthesis, N-glycan biosynthesis, sphingolipid metabolism, glycosphingolipid biosynthesis, pyruvate metabolism, xenobiotic metabolism by cytochrome P450, pyrimidine metabolism, and glycerolipid metabolism (sex differences can be found in Supplementary Figure S4). These pathways are linked to cellular signalling^46–48^, matrix composition^49,50^, matrix remodeling^48^, cell-matrix interactions^46^, stress response^47,49^, and cell viability^51^.

Because not all groups of the HFD and aging signatures had the same degree of cellular and matrix alterations, we also determined group-specific differences in metabolism. ECCO was used in specific two-group comparisons (e.g., young HFD females vs aged HFD females) to identify statistically significant pathways (gamma p < 0.05) for each group (Figure 9c). HFD in young females was associated with upregulation in amino acid metabolism, compared to LFD young females and young HFD males. Young HFD males, the group with the highest adiposity, also had downregulated amino acid synthesis compared to young LFD males. Aged HFD females showed an attenuated amino acid metabolism compared to aged HFD males and young HFD females but had no significant differences to aged LFD females. Aged HFD males had upregulated amino acid metabolism compared to young HFD males, but there was no difference to aged LFD males. Aged LFD mice had no significant sex differences (Figure 9d).

Inflammatory metabolic pathways, such as butanoate, histidine, and nicotinamide pathways, were upregulated in young HFD females compared to young LFD females. Aged HFD females exhibited minor downregulation of these inflammatory pathways compared to young HFD females. Aged HFD males demonstrated upregulation of the same inflammatory pathways compared to their young counterparts and aged females. Young HFD males downregulated these pathways compared to young LFD males and females of both diets, while upregulating other inflammatory pathways, such as the folate-mediated one carbon pool. Differences in the HFD response between young females and males may reflect sex differences or variation in adipose gains.

The few reports of cortical bone metabolomics have identified sex differences^52,53^, but this area of investigation would benefit from additional datasets. Therefore, we compared sex differences in this study with those from a previous study by our group on 5-month-old male and female C57BL/6JN mice, which were bred at Montana State University and fed standard chow diet^53^. Welhaven et al. used the same method of metabolite extraction reported here followed by MetaboAnalyst analysis. We analyzed cortical bone tissue metabolism from young LFD females versus males using MetaboAnalyst and ECCO. Both studies agreed that young LFD male mice have a more pronounced signature of amino acid metabolism compared to young LFD female mice (Figure 9d). However, there was less agreement on lipid metabolism; the prior study identified upregulated fatty acid metabolism in females, whereas the present study did not detect this sex difference, possibly due to dietary differences (i.e., standard chow versus low-fat diet).

In summary, these data demonstrate that the distinct cortical bone metabolic signature for HFD is associated with pathways related to glucoregulation. By contrast, the distinct aging metabolic signature is associated with pathways related to cellular function decline and death. Both HFD and aging metabolic signatures share pathways related to matrix signaling and regulation, which aids in understanding the loss of bone fracture toughness seen in both groups.

## 3.0 Discussion

This study tested the hypothesis that HFD exacerbates the loss of bone toughness in aging but that aging and HFD differently dysregulate bone matrix properties and cortical bone tissue metabolism. Mice were fed 45% HFD for 8 weeks, which resulted in varying degrees of adiposity within and between experimental groups. However, glucose metabolism was disrupted for all HFD groups. We found that aging and HFD each reduced bone toughness in an additive fashion, but likely for different reasons. Aging primarily impacted bone tissue characteristics related to mineral maturity, while HFD mineral content and maturity as well as collagen structure. Cortical tissue metabolism demonstrated both overlapping and distinct signatures from aging and HFD. These findings suggest that dietary fat, regardless of adipose accumulation, can worsen the loss of bone fracture resistance in aging for both sexes.

Our study demonstrates that the causes of whole-bone fragility in aging and HFD have important differences. Aging reduced bone strength and fracture toughness, while HFD only reduced fracture toughness but not strength. Both aging and HFD reduced the stress intensity factor at crack initiation (Kc_init_), which measures the intrinsic toughness of bone matrix (i.e., mechanisms that act in front of the crack tip)^54^, but HFD did not lower the stress intensity factor at the maximum load (Kc_max_), which includes both intrinsic and extrinsic toughening mechanisms (i.e., mechanisms that act in front of the crack tip and behind the crack’s wake)^54^. Thus, aging and HFD likely impact essential bone matrix properties promoting intrinsic toughening mechanisms in both sexes but not extrinsic mechanisms. Future work should attempt to identify specific intrinsic toughening mechanisms impacted by aging and HFD. Importantly, the additive effects of aging and HFD on decreasing bone toughness indicate that the negative impact of HFD on bone toughness is particularly detrimental in the context of aging.

Despite similar whole-bone toughness outcomes in both sexes, the effects of HFD and aging on matrix properties differed between sexes. Two regions of the cortical femur were selected for hydrated Raman spectroscopy analysis of matrix properties: the periosteal surface and an endocortical region of the femur cross-section. At the periosteal surface, age effects on matrix properties were more apparent than HFD. These measurements recapitulated several changes in aging rodent^55^ and human bone^56^ that have been seen in prior work, such as increased mineral carbonate content and crystallinity. At the endocortical surface, the effect of HFD on bone matrix became apparent, but in a manner that interacted with age and sex. For young females, HFD increased bone mineral:matrix ratio and also increased the spatial gradient of mineral maturity, as seen in the measures of carbonate:phosphate and crystallinity. Females also had decreased spatial variation in collagen secondary structure in response to HFD. These changes were not seen for males, indicating sex differences in the impact of age and diet on bone matrix properties, despite reductions in whole bone toughness in both sexes. The specific reasons for reduced bone toughness with age and diet do not appear to be conserved across sexes at the matrix level, underscoring the need to study sex differences in the impacts of aging on bone biomechanics.

Because measures of bone matrix quality have the potential to intercorrelate, we tested the association of fracture toughness and bone matrix properties using an unbiased clustering approach^40^. In this approach, measures with the strongest absolute correlations are clustered together. At the periosteal surface, the cluster with the strongest correlations with fracture toughness included collagen secondary structure (I1670/I1640) and carbonate:phosphate. At the endocortical surface, the cluster with the strongest correlations with fracture toughness included collagen secondary structure (I1670/I1610), variation of collagen secondary structure (RMSD 1670/I1610), and mineral:matrix ratio. Indicators of mineral maturity, such as carbonate:phosphate and crystallinity, and fAGEs were less associated with fracture toughness. These data align with studies demonstrating stronger associations between fracture toughness and collagen integrity in human^56,57^, bovine^58^, and rodent^55^ bones, and weaker associations with mineral maturity^59^. Importantly, our data reveal insights into how toughness and matrix properties cluster by age and diet at different locations. For example, at the periosteal surface, data cluster by age more than by diet. However, at the endocortical surface, data cluster in a manner that suggests important impacts from both age and diet. These data also challenge the interpretation of how fAGE accumulation impacts bone fracture resistance. fAGE accumulation has been shown to have significant associations with decreased bone toughness in the contexts of diabetes and kidney disease, which is thought to be related to AGE crosslinks hindering intrinsic toughening mechanisms^39,60^. However, our data do not show an association between fAGE accumulation and reduced fracture toughness in aging, suggesting that greater perturbations in fAGE accumulation than produced in the current study may be needed to reduce fracture toughness. Another possible explanation may be that fAGE-related signaling rather than the accumulation is a more important contributing factor to reduced fracture toughness in HFD and aging. This is supported by a study by Stephen et al. where AGE receptor knockout mice had improved elasticity:hardness ratios, which is a proxy measure for fracture toughness, in young adult female mice regardless of diet and low-fat diet young adult males^61^. Chronic elevated AGE signaling may also contribute to bone fragility conditions in aging humans^62^.

Metabolomic assessments of cortical bone tissue can reveal connections between altered bone matrix properties and the cellular response to aging and HFD. Aging and HFD groups exhibited distinct metabolic signatures largely independent of sex. For example, HFD groups had upregulated branched chain amino acid biosynthesis, while aging groups had downregulated pentose phosphate pathway. The distinct response associated with HFD groups was the progression of glucose dysregulation^41^, whereas aging groups demonstrated decreased cellular functionality^43^, increased stress response^42^, and increased cell death^45^. Both aging and HFD signatures shared overlapping pathways, including GPI-anchor, N-glycan, and sphingolipid biosynthesis. These pathways are associated with matrix regulation^48^, stress responses^47,49^, turnover activity^48^, and cell viability^51^. It is also notable that some of these pathways can be downstream of AGE signaling^47^. Further studies are required to determine the role of these metabolic pathways in matrix regulation and bone fragility. These data illustrate the distinct and shared bioenergetics of aging and HFD in bone fragility.

Adipose-induced inflammation can influence bone cell metabolism, glucose tolerance, and regulation of bone matrix properties^20,33^. Although neither HFD nor aging promoted a clearly elevated systemic inflammatory response, metabolites from cortical bone tissue showed that all HFD and aging groups had local inflammatory responses. Young females on HFD demonstrated resistance to the inflammatory environment through upregulation of histidine metabolism and branched chain amino acid degradation, which are necessary to attenuate oxidative stress and inflammation^63,64^. Increased biosynthetic pathways related to amino acids were also associated with this adaptivity towards HFD. Aged HFD females and males had similar inflammatory and amino acid synthesis pathways as the young HFD females but with lower activity. In contrast, young HFD males, the group with the highest adiposity, exhibited a markedly different inflammatory response compared to other HFD groups. The response of young HFD males was characterized by lower histidine metabolism and higher folate-mediated one carbon pool pathways. The one carbon pool by folate synthesizes amino acids important for cellular growth and is essential for maintaining oxidative balance within the cell^65^. The general patterns of inflammatory responses of cortical bone align with studies of aging and HFD in other mouse tissues, particularly adipose tissue^13,14^. These data suggest that HFD induces sex- and age-specific responses to local inflammation, potentially contributing to variations in bone matrix regulation.

HFD likely has a combination of direct and indirect effects on the regulation of bone matrix properties. For example, sphingolipid metabolism may have a direct effect on bone matrix since accumulation of this pathway’s byproducts can hinder osteoblast activity and regulate apoptosis^48^. In contrast, linoleic acid could indirectly affect bone matrix since it may alter peroxisome proliferation signaling, which can regulate osteoblast activity^66^. Leptin also indirectly regulates bone matrix turnover, through one or more mechanisms^67^. Further mechanistic studies will improve knowledge about how dietary fat can impact bone cells and their regulation of bone matrix.

Osteocytes are the main regulators of bone matrix, indirectly through surface bone turnover and possibly directly through lacunar-canicular system (LCS) turnover^68^. We investigated whether aging and HFD affect these processes. We found that osteocyte viability was decreased with HFD, but only in aged mice, indicating that the interactive effects of HFD and age compromise long-term bone quality regulation by osteocytes. Aging also altered LCS turnover, evidenced by lower *Mmp2* and higher *Acp5* gene expression. HFD upregulated vacuolar ATPase gene expression, suggesting increased LCS resorption. This study is not comprehensive in assessing osteocytes under HFD, and other studies indicate deleterious effects of HFD on *in vitro*^17^ and *in vivo*^69^ osteocyte function. Further mechanistic inquiry is needed to understand the role of the osteocyte in maintaining bone quality in aging with HFD in both sexes.

This study has several limitations. Several measures may have been underpowered to detect effects of age and HFD (e.g., cortical thickness and several measures from Raman spectroscopy). In addition to the variability of aging on bone outcomes^70^, both age groups had responders and non-responders to HFD, contributing to variability in bone measurements. The sex differences identified in young female versus male mice may reflect differences in adiposity rather than true sex differences. To address this challenge, we used the AUC from glucose tolerance tests as a covariate in statistical models. Furthermore, adipose tissues can directly or indirectly signal to bone^24^, but these responses were not investigated in this study and should be explored in future investigations. Raman spectroscopy was only performed on half of the female samples due to sample availability, limiting our ability to estimate sex differences in matrix effects. The fAGE analysis of humeri may not accurately reflect fAGE accumulation in femur tissue and does not measure non-fluorescent AGEs, which may influence bone mechanical properties^39^. While we measured many features of matrix composition and maturity, additional measurements related to collagen crosslinking and non-collagenous proteins would be valuable in future studies.

In conclusion, HFD has detrimental effects on skeletal fracture resistance in both young adult and early-old-age C57BL/6JN mice. Regardless of adiposity and inflammatory response, HFD decreased fracture toughness for females and males. HFD and aging both deleteriously impacted bone matrix, but in distinct ways that depended on sex. Further, there were shared and distinct aspects of the cortical bone tissue metabolic dysregulation induced by HFD and aging, which suggests that the loss of fracture resistance produced by each factor is produced as a consequence to different changes to bone cells and their surrounding matrix. Together, these results demonstrate that short-term HFD has important negative effects on bone fracture resistance and that HFD has the potential to exacerbate the bone fragility of aging.

## 4.0 Methods

### 4.1 Animal and tissue harvest

All animal procedures were approved by Montana State University’s Institutional Animal Care and Use Committee. Female and male C57BL/6JN mice (National Institute of Aging Aged Rodent Colony at Charles River Laboratories, Wilmington, MA) were housed in cages of 2-5 mice with chow and water *ad libitum* on a 12-hour light-dark cycle. Mice were randomly assigned to either an 8-week high-fat diet (HFD; Research Diets, D12451, 45% fat) or low-fat diet (LFD; Research Diets, D12450H, 10% fat) such that there were 10-11 mice/group. Mice were weighed weekly. Calcein (20 mg/kg) and alizarin (30 mg/kg) were respectively administered via intraperitoneal injection at eight and two days before euthanasia.

Mice were overdosed with isoflurane and exsanguinated via cardiac puncture at ages 5-(Female LFD: n = 10; Female HFD: n = 10; Male LFD: n = 10; Male HFD: n = 11) and 22-months (Female LFD: n = 9; Female HFD: n = 9; Male LFD: n = 8; Male HFD: n = 9).

Several aged mice from both diet groups did not reach the study endpoint and were excluded due to tumors or other gross pathologies.

Serum was collected from the cardiac puncture and stored at -80°C. Both femora were stored in phosphate-buffered saline (PBS)-soaked gauze without soft tissues and frozen at - 20°C for later mechanical testing. The left tibiae were flushed of marrow with PBS. The marrow and flushed cortical bone were stored separately at -80°C. Perigonadal fat pads were weighed as a surrogate measure of adiposity.

### 4.2 Intraperitoneal glucose tolerance test (ipGTT)

ipGTT was performed seven days before euthanasia. Mice were fasted for six hours following the dark cycle. Blood glucose was measured at baseline (0 min), 15, 30, 60, 90, and 120 minutes after glucose administration (2 g dextrose per kilogram body mass injected into the intraperitoneal cavity) from the tip of their tails using a blood glucometer (Accu-check Performa, Roche, Indianapolis, IN, USA). Baseline glucose, maximum glucose, delta glucose (maximum-baseline glucose), and glucose integrated area under the curve (glucose AUC) were calculated.

### 4.3 Trabecular microarchitecture and cortical geometry

Left femurs were assessed for microarchitecture and geometry using a high-resolution micro-tomographic imaging system (µCT40, Scanco Medical AG, BrulJttisellen, Switzerland). Image acquisition and analysis followed JBMR guidelines^71^ with scanning parameters of isotropic voxel size of 10 µm^3^, peak X-ray tube potential of 70 kV, tube current of 114 µA, 200 ms integration time, and Gaussian filtration was applied to scans. The distal metaphysis was assessed for trabecular microstructure beginning 200 µm above the distal growth plate to 1500 µm proximal. The trabecular region was identified by manual contouring of the endocortical region with thresholding at 375 mgHA/cm^3^. Scanco Trabecular Bone Morphometry script was then used to measure trabecular parameters. Cortical geometry was assessed in 50 transverse sections of the cortex area along a 500 µm long region of the mid-diaphysis for cortical geometry. Cortical bone to calculate cortical parameters from was identified using a threshold of 700 mgHA/cm^3^. Femurs were kept in PBS-saturated gauze at –20 °C before and after testing.

### 4.4 Biomechanical testing of whole-bone properties

All biomechanical tests utilized an Instron 5543 (1 kN load cell, Instron, Norwood, MA). Material properties of left femora were evaluated using three-point bending. PBS-hydrated femora were tested until failure at a rate of 5 mm/min on a custom fixture with an 8 mm span. Load-displacement data and micro-computed tomography measures of cross-sectional geometry were used to calculate the modulus, ultimate compressive strength, and toughness as calculated from area under the curve until fracture using standard bone flexural testing equations^72^.

Notched fracture toughness was evaluated on right femora consistent with Welhaven et al.^53^. Briefly, femurs were notched at the posterior surface of the mid-shaft to a target notch depth of 1/3 of the anterior-posterior width. Notched femora were loaded with the notch facing down in three-point bending until failure at a loading rate of 0.001 mm/sec. Distal halves of fractured femurs were flushed of marrow after testing, air dried, and imaged using variable pressure scanning electron microscopy (20 Pa, 15 kV; Zeiss SUPRA 55VP, Oberkochen, Germany). Cortical bone geometry and the initial notch half-angle were obtained by analyzing the images of the fracture surfaces using a custom MATLAB code. Fracture toughness values were calculated for two criteria. The Kc_max_ measure calculates the stress intensity factor at the maximum load given the starting notch geometry. The Kc_init_ measure calculates the stress intensity factor when the crack begins to propagate, using the load at yield given the starting notch geometry. These measures and their interpretation are described in Ritchie and co-authors^73^.

### 4.5 Quantitative reverse transcription polymerase chain reaction (qRT-PCR)

Flushed left tibiae were processed according to a modified version of the procedure by Kelly et al^74^. Cortical bones were powdered in liquid nitrogen. RNA was isolated using Trizol (Sigma-Aldrich, St. Louis, MO) and RNeasy Mini Kits (Qiagen, Hilden, Germany). Reverse transcription was done using High-Capacity cDNA kits (Applied BioSystems, Waltham, MA) and qRT-PCR analyses using PowerUp SYBR green master mix (Applied BioSystems, Waltham, MA) on an Applied Biosystems Quant Studio 5, (Waltham, MA). All kits were used according to manufacturer’s protocol. Primer sequences are reported in Supplementary Table S12.

### 4.6 Serum biomarkers

Enzyme-linked immunosorbent assay (ELISA) kits were used according to manufacturer protocols on blood serum collected at tissue harvest. Measurements included CTX1 (Mouse Cross Linked C-telopeptide of Type I collagen Mini Samples ELISA kit, MyBioSource, San Diego, CA), P1NP ( Mouse procollagen I N-terminal peptide, P1NP ELISA kit (MyBioSource, San Diego, CA), adiponectin (Quantikine ELISA Mouse Adiponectin/Acrp30 kit, R&D Systems, Minneapolis, MN), resistin (Quantikine ELISA Mouse Resistin kit (R&D Systems, Minneapolis, MN), and interleukins 4, 6, and 10 (IL-4, IL-6, IL-10), fibroblast growth factor 21 (FGF21), leptin, vascular endothelial growth factor (VEGF), C-reactive protein (CRP), and complement factor D/adipsin (CFD/adipsin) (Luminex Discovery Assay Mouse Premixed Multi-Analyte kits, R&D Systems, Minneapolis, MN).

### 4.7 Metabolomics

Metabolomic profiling was performed following methods established by our groups using flushed right humeri bone^53^. Briefly, metabolites from pulverized bone were precipitated with 3:1 methanol:acetone, vacuum concentrated, and resuspended in 1:1 acetonitrile:water before analysis using untargeted liquid chromatography mass spectrometry (Agilent 1290 LC coupled with Agilent 6538 Quadrupole-Time of Flight (Q-TOF) mass spectrophotometer in positive mode, Cogent Diamond Hydride HIILIC column).

Raw spectra of detected metabolites were aligned, peak picked, and processed using MSCConvert and XCMS. Metabolite data was then analyzed using ensemble clustering with cluster optimization (ECCO)^40^. ECCO utilizes unbiased methods to 1) determine the optimal number of clusters required for the analysis of a given dataset (e.g. K-means, Dunn, PBM indexes) and 2) finds 13 clustering solutions through distance algorithms, where each metabolite cluster is then weighted. The sum of these weights across all cluster solutions is then represented as clusters of metabolites grouped by frequency that they group together with 1.0 meaning the metabolites are together in 100% of the solutions^40^. The contents of each cluster were assessed using Mummichog in MetaboAnalyst to find significant (gamma p < 0.05) or non-significant pathways (gamma p ≥ 0.05). Filtering of the pathways was performed for non-significant pathways to reduce duplicates and pathways that were significantly different in at least one other metabolic statistical comparison between groups. The resulting collection of non-significant pathways were called signatures. Signatures were compared to their respective control (e.g. HFD metabolites in the signature pathways compared to LFD values).

Since aging and HFD each decreased fracture toughness in this study, ECCO-based analyses were also used to investigate shared and distinct metabolic contributions. The metabolic signatures of aging and HFD were generated. The metabolic pathways identified with in each factor’s signature were compared to other factor signatures. ‘Shared’ pathways were those that were found in multiple signatures (i.e. both aging and HFD). ‘Distinct’ pathways were found in only one signature (i.e. only aging or HFD, Supplementary Figure S5).

To determine the metabolic differences between specific two-group comparisons, ECCO was used to identify statistically significant pathways (gamma p < 0.05) within clusters. 2-group comparisons were performed for all relevant age, diet, and sex comparisons (Figure 9c).

### 4.8 Histology

Right tibiae were decalcified with EDTA disodium salt dihydrate, dehydrated in a graded ethanol series, embedded in paraffin, and serially sliced into 5-micron thick longitudinal sections. Slices of each tibia were stained with either Terminal deoxynucleotidyl transferase dUTP Nick End Labelling (TUNEL), Tartrate resistant acid phosphatase (TRAP), or Hemotoxin and Eosin stain (H&E). The proximal tibial metaphysis was imaged using a Nikon E800 (4x and 10x). TUNEL was used to determine percent apoptotic osteocytes and osteocyte density per area. TRAP was used to determine osteocyte density per metaphyseal medullary cavity perimeter. H&E was used to identify bone marrow adipocytes using a custom MATLAB code. Mean adipocyte area, marrow cavity area, and adipocyte number density were reported.

Quantitative histomorphometry was performed on diaphyseal cross sections of polished embedded femurs to determine local turnover using confocal laser scanning microscopy (5x magnification, Leica SP5, Wetzlar, Germany) with emission ranges of 490-550 nm for calcein and 580-680 for alizarin. The percentage of mineralizing bone surface (MS) was calculated for each label for the endocortical (E. BS), periosteal (P. BS), and total surfaces (BS).

### 4.9 Periosteal and cross-sectional hydrated Raman spectroscopy

Following three-point bending, Raman spectroscopy was performed for the femoral midshaft at two locations, the periosteal surface and the femur midshaft cross-section. The periosteal surface was chosen because the bone was minimally processed (i.e., no cutting or polishing) to achieve these measurements. The femur cross-section was also investigated because an endocortical mineralization defect in response to HFD was detected in other study analyses. In both cases, bones were studied under hydrated conditions.

For the periosteal surface, data was collected at five equidistant points (500 µm apart) along the posterior shaft of the distal end of hydrated left femora using a 10x dry objective lens. The bones were not embedded. Samples were kept hydrated using a custom tap water-saturated sponge mount. Raman spectra were collected with a frequency range of 300-1900 cm^-1^ (Horiba Laser Tweezer Raman Modified LabRam HR Evolution NIR, Kyoto, Japan) using a 785 nm laser. A 12^th^ order polynomial baseline correction was applied to spectra using LabSpec6 software. Area ratios were calculated for mineral:matrix (v_2_PO_4_/Amide III) and carbonate:phosphate (v_1_CO_3_/v_1_PO_4_). The inverse of the width of the of the v_1_PO_4_ peak at half of the maximum intensity (1/FWHM) was used to calculate the crystallinity^75^. Subpeaks of Amide I at 1610, 1640, and 1670 cm^-1^ were determined through the second derivative method^75^. Intensity ratios included I1670/I1640 and I1670/I1610, which are sensitive to collagen secondary structure^75^. Custom MATLAB code was used to calculate all measures. For each sample, the median of the five points was input into statistical models to avoid overpowering analyses.

For the endocortical cross-sectional map, the same distal femoral ends were embedded in non-infiltrating viscous epoxy (EpoxiCure 2, Buehler, Lake Bluff, IL). A cross section was cut at the mid-diaphysis and polished using a graded alumina series. Ethanol was avoided during the process. Cross-sections were stored at -20 °C prior to analyses and then rehydrated with tap water for 4 hours before Raman measurements. Maps were placed in the anterior-lateral region of the cortical section, starting at the endocortical surface (3x10 with points 20 µm apart in x-direction and 7 µm apart in the radial direction) using a 60x water immersion lens. Samples were kept hydrated during testing by the column of tap water between the lens and sample. Points with an epoxy/ v_1_PO_4_ ratio greater than 7.5% were discarded as these points were contaminated with plastic. For other points, epoxy was subtracted from the baseline, although the epoxy signal was usually minimal or negligible. A custom Python code (Version 3.9) was used to calculate the normalized position of points relative to the cortical thickness. Mineral:matrix, carbonate:phosphate, crystallinity, and amide I subpeak measurements were calculated as described above. The median measurements of the measures, as well as their slopes (gradient) and root mean square deviation (RMSD) were calculated for 0-30% of the cortical thickness to determine the change of these measures with distance from the endocortical bone-forming surface. 30% was chosen as the maximum distance because this distance was measured for all study mice (Supplementary Figure S5).

### 4.10 Cortical bone matrix fluorescent AGEs (fAGEs)

fAGEs were quantified from the diaphysis of flushed left humeri using previously described protocols^76^. Briefly, the fat was removed from samples by three 15-minute washes with agitation in 200 µL of 100% isopropyl ether. Then samples were lyophilized for 8 hours (FreeZone 2.5 L freeze-dry system, Labconco, Kansas City, MO) before hydrolyzation for 20 hours at 110°C in 6 N HCl (10 µL/mg sample). The solutions were diluted 1:100 followed by centrifugation at 4°C and 13000 rpm. Dilutions were stored at -80 °C in dark conditions. Quinine fluorescence was measured at 360/460 nm excitation/emission for both sample and stock solutions (stock 10000 ng/mL quinine sulfate / 0.1 N H_2_SO_4_, Biotek Synergy HTX Multi-mode Reader, Winooski, VG). Separately, hydroxyproline was oxidized in both sample and stock solutions (stock 2000 µg/mL L-hydroxyproline / 0.001 N HCl) with chloramine-T solution for 20 minutes. Residual chloramine-T was quenched with perchloric acid (3.15 M) and incubated for 5 minutes at room temperature. Solution was incubated at 60°C for 20 minutes after p-dimethylaminobenzaldehyde was added. The solutions were kept in dark conditions while cooling to room temperature. The same plate reader measured the absorbance at 570 nm to quantify collagen from hydroxyproline^76^, which was used to normalize fAGE content for reporting (ng quinine/mg collagen).

### 4.11 Statistics and reproducibility

Three-way ANOVA tests were performed to determine if bone outcomes depended on age, sex, diet, or interactions of these factors (Minitab, v.21.2). Response variables were transformed if required to satisfy model assumptions of residual normality. The criterion for significance was set to *a priori* to p < 0.05. Significant interactions were followed up with post-hoc tests conducted using Fisher pairwise comparisons corrected using the Bonferroni procedure to control family wise error (i.e., the criterion for significance is adjusted to 0.05 / number of comparisons). ANCOVA determined if glucose AUC was a significant covariate for all models. When covariates were not significant, the model was rerun without the covariates to interpret main effects and interactions. Analyses was performed with and without outliers identified by Grubb’s Test. Results are presented without outliers, but the majority of conclusions made from statistical analyses did not differ whether outliers were included or not. The 11 measures that did not have the exact same conclusions with and without outliers are presented in Supplementary Table S13. Specific sample n for each ANOVA or ANCOVA test can be found in the provided Supplementary Tables S1-S11.

To generate the clustered heatmaps, the group medians of the Raman, fAGE, and fracture toughness (Kc_init_) results were normalized and scaled by feature using min-max normalization. Scaling by feature indicates that each variable for the heatmap was scaled to have values 0-1, so that all the variables had comparable ranges. Missing values were removed. ECCO utilized correlation metric distances (Equations 1 and 2) to reach a consensus of the features in the optimal number of clusters required to describe the data. The correlation methods weigh negative and positive correlations equally, so that clusters can have inverse relationships within them. Each cluster represents which features were most strongly correlated to each other.

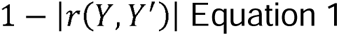

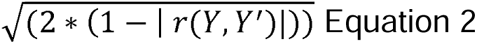

Here, *r*(*Y*, *Y*’ ) is the correlation distance between two given data points *Y* and *Y*’. The resulting correlation coefficients were condensed using the squareform package in Python, which returns a symmetric distance matrix. Then, the coefficients were input into the linkage function with the average linkage method. Within ECCO, the resulting clusters and linkages were then used to create a cluster heatmap of the raw data using Python’s Seaborn library clustermap function.

## Supplementary Information

All supplementary tables and figures can be found in the supplementary files.

## Data Availability

We will provide data in response to reasonable requests.

## Code Availability

ECCO can be found at https://github.com/hisl6802/ECCO.

Code used for biomechanical and Raman analyses is available from the investigators upon reasonable request.

## Acknowledgements

We thank Maria Jerome for preparing histology samples, Dr. Heidi Smith from the Center for Biofilm Engineering for assisting with microscopy, The Center for Advanced Orthopedic Studies at Beth Israel Deaconess Medical Center for performing the micro-tomographic imaging, and Dr. Albert Parker for advice on statistical analyses.

This research was made possible by the Department of Mechanical & Industrial Engineering and the College of Engineering at Montana State University. We are grateful for support from the National Science Foundation (2120239, 2340823) and the National Institute of Health (AG068680). This work represents the views of the authors and not necessarily those of the sponsors.

## Author Contributions

**K.B.** Methodology, Software, Investigation, Formal Analysis, Data Curation, Writing – Original Draft, Writing – Review & Editing

**G.V.** Investigation, Formal Analysis, Writing – Original Draft, Writing – Review & Editing

**B.D.H.** Methodology, Software, Formal Analysis, Writing – Review & Editing

**M.M.** Investigation, Formal Analysis, Writing – Original Draft, Writing – Review & Editing

**S.W.** Methodology, Investigation, Writing – Review & Editing

**H.D.W.** Methodology, Investigation, Writing – Review & Editing

**R.B.** Investigation, Writing – Review & Editing

**K.O.P.** Investigation, Writing – Review & Editing

**L.K.** Formal Analysis, Resources, Writing – Review & Editing

**R.K.J.** Formal Analysis, Resources, Writing – Review & Editing

**S.A.M.** Conceptualization, Formal Analysis, Resources, Writing – Original Draft, Writing – Review & Editing

**C.M.H.** Conceptualization, Formal Analysis, Resources, Writing – Original Draft, Writing – Review & Editing

