## Supplementary figures and tables for "High-fat diet exacerbates the loss of bone fracture toughness in aging for C57BL/6JN mice"

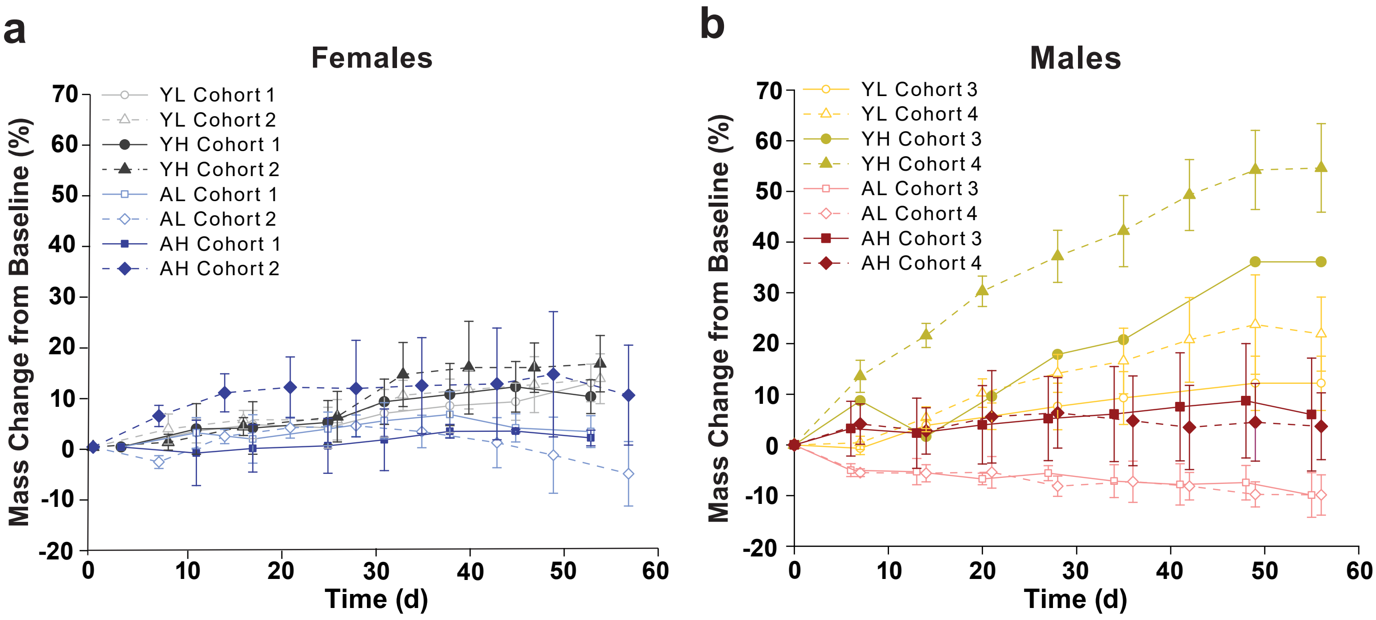


**Figure S1. Female cohort mass gain trajectories over the course of the diet intervention were less variable than male cohorts on high fat diet.** Mass gain trajectories for the cohorts during the diet intervention of (a) female and (b) male C57BL/6JN mice. Line plots are presented as mean ± 1 SD. YL = Young (5-month) low-fat diet, YH = Young (5-month) high-fat diet, AL = Aged (22-month) low-fat diet, AH = Aged (22-month) high-fat diet.

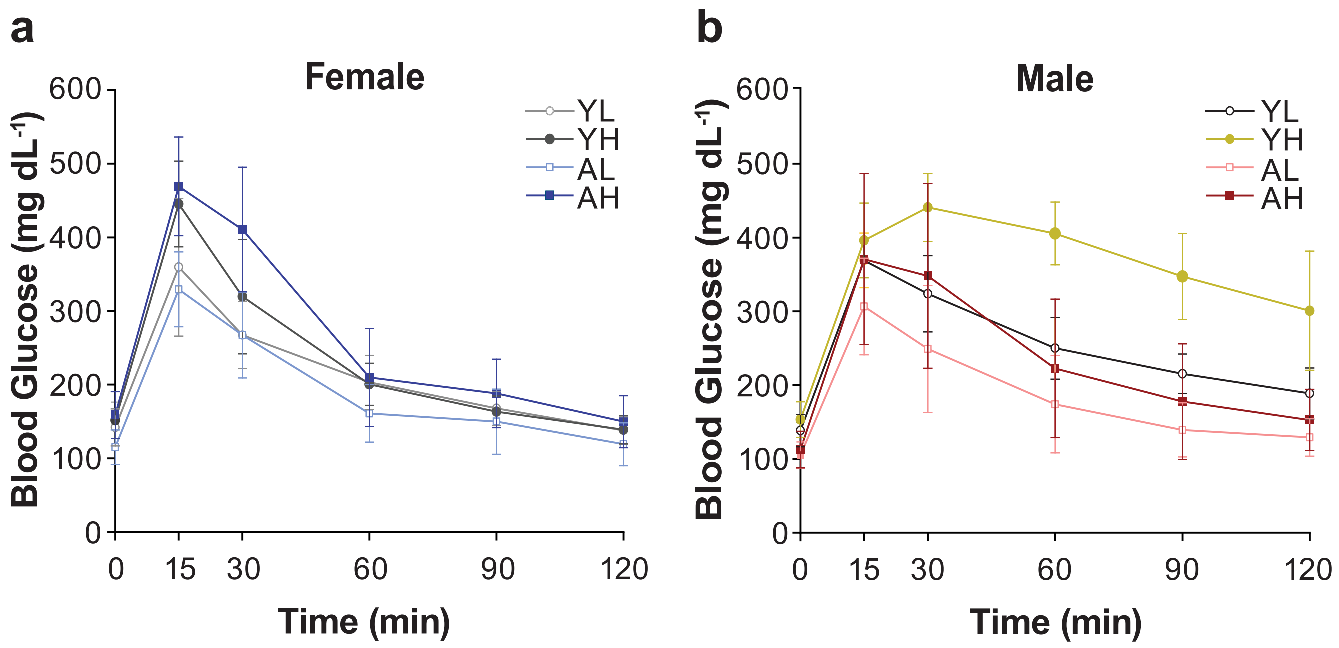


**Figure S2. All high fat diet groups had elevated baseline, delta, and maximum glucose values regardless of adiposity developed.** Intraperitoneal glucose tolerance testing (ipGTT) test curves for (a) female and (b) male groups. Line plots are presented as mean ± 1 SD. YL = Young (5-month) low-fat diet, YH = Young (5-month) high-fat diet, AL = Aged (22-month) low-fat diet, AH = Aged (22-month) high-fat diet.

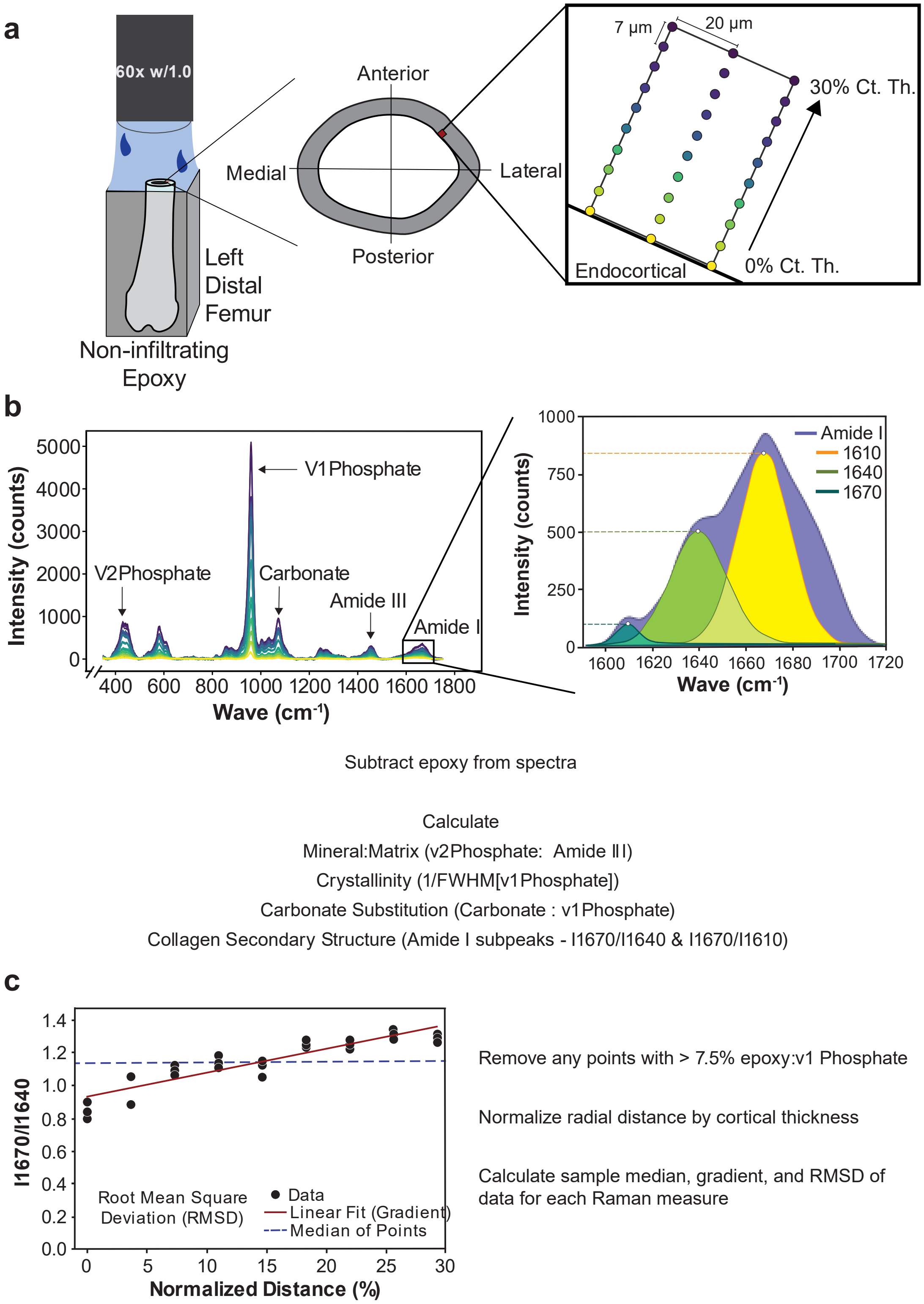


**Figure S3. Steps for hydrated cross-sectional Raman spectroscopy.** (a) The general setup for left distal femurs to map the anterior-lateral region of interest in hydrated conditions. Each point in the mapping region had (b) a corresponding spectrum. The epoxy signal was subtracted from each spectra before the five measures were calculated. The amide I peak was analyzed for subpeaks for 1610, 1640, and 1670 cm$^{-1}$. The mineral:matrix ratio was calculated by the ratio of the $v_2$ phosphate and amide III peaks, crystallinity was calculated with the inverse of the full width half maximum intensity of the $v_1$ phosphate peak, carbonate substitutions in the mineral was calculated as the ratio of the carbonate and $v_1$ phosphate peak, collagen secondary structure ratios were estimated as the intensities ratios of the amide I subpeaks I1670/I1610 and

I1670/I1640. (c) The measures were plotted with the normalized distance to find a linear fit. The slope of the linear fit was used as the gradient for a measure. The gradient represents the slope of change for a measure with respect to the radial distance from the endocortical surface. The root mean square deviation (RMSD) was calculated from the residual sum of squares about the linear fit and the number of data points in the 0-30% region. The RMSD is an estimate for the spatial variance of a measure. The median of all the points in the 0-30% region was used as the median.

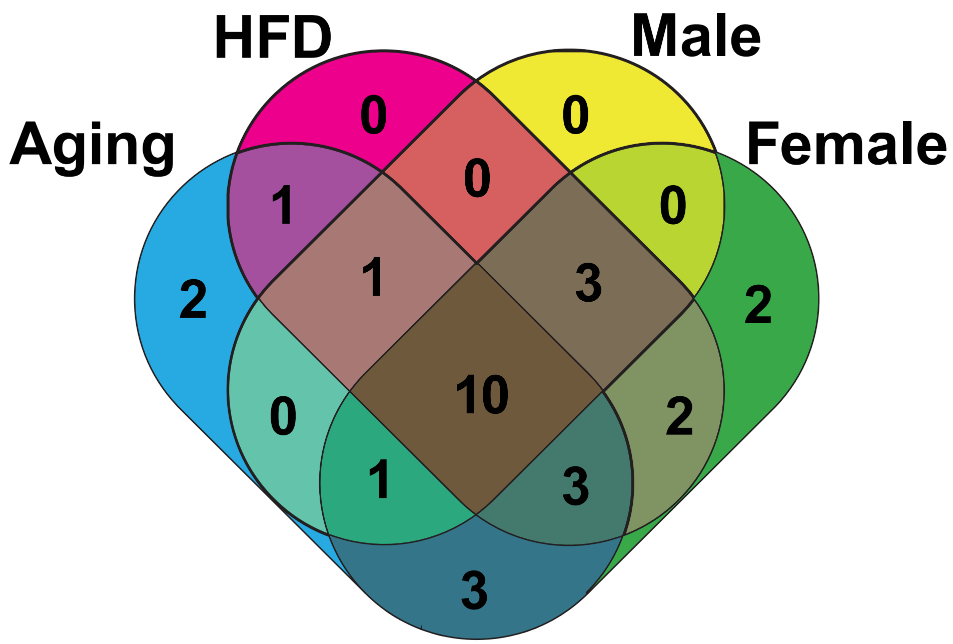


| | |
|---|---|
| HFD<br>Aging | Tryptophan Metabolism |
| Male<br>HFD<br>Aging | Steroid Hormone Synthesis |
| Female<br>HFD<br>Aging | Pyruvate Metabolism<br>Beta-Alanine Metabolism<br>Metabolism of Xenobiotics by Cytochrome P450 |
| Male<br>Female<br>HFD<br>Aging | N-Glycan Biosynthesis<br>GPI-Anchor Biosynthesis<br>Glycosphingolipid Biosynthesis<br>Sphingolipid Metabolism<br>Alpha-Lenolenic Acid Metabolism<br>Arachidonic Acid Metabolism<br>Glycerolipid Metabolism<br>Pyrimidine Metabolism<br>Purine Metabolism<br>Cysteine and Methionine Metabolism |

**Figure S4. High-fat diet and aging metabolic signatures are shared between female and male signatures.** A summary of shared and distinct metabolic pathways for high-fat diet (HFD), aging, male, and female signatures.

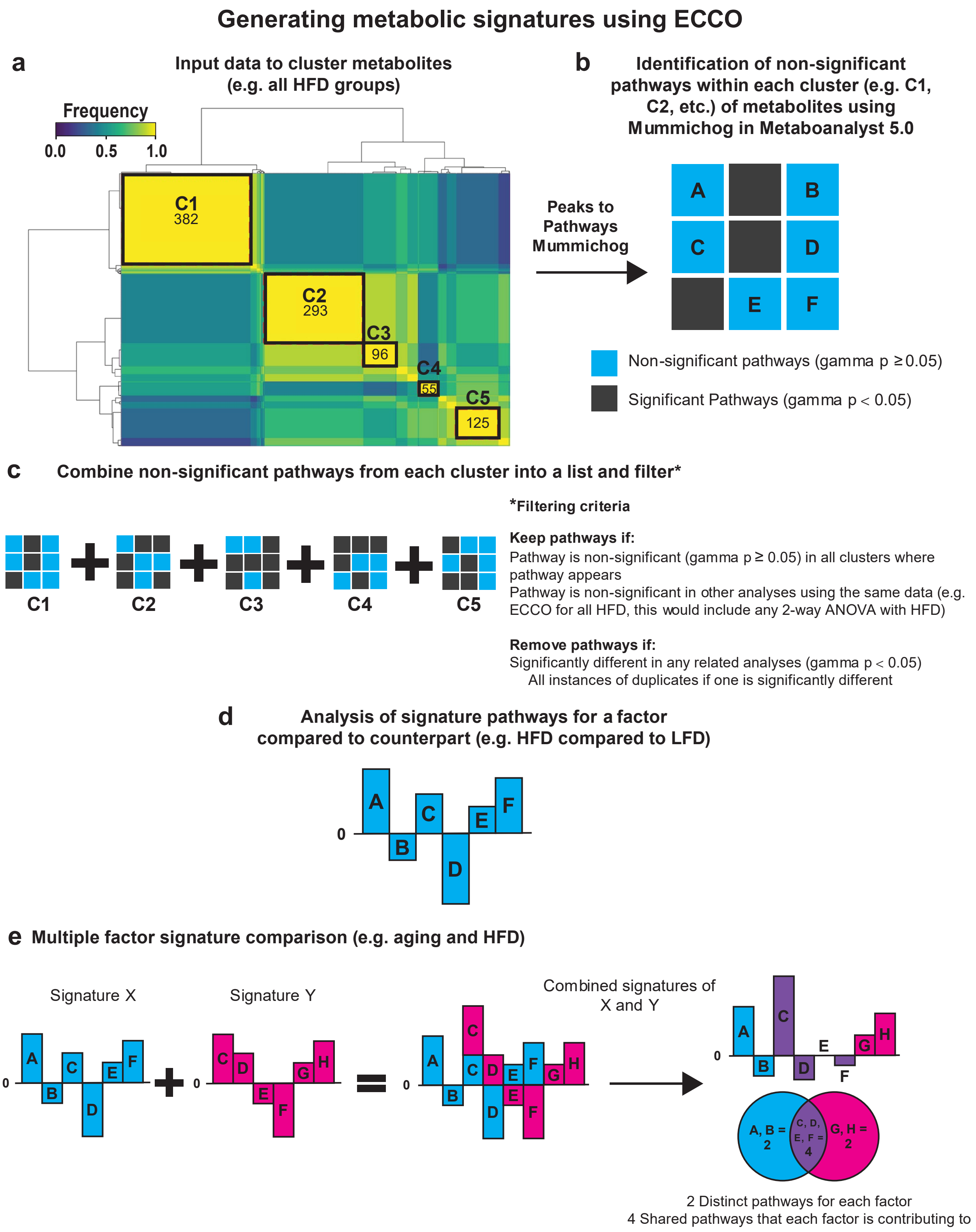


**Figure S5. Steps for assessing metabolomic data using ensemble clustering with cluster optimization (ECCO) to identify signatures for multiple factors.** (a) Metabolites are clustered according to the frequency the metabolites group together in multiple clustering solutions. (b) Clustered metabolites can be identified using Mummichog. (c) The resulting pathways are grouped into non-significant or significant according to the threshold for the gamma value. The pathways from the clusters are filtered to avoid duplicates and significant pathways from other analyses. (d) The remaining signature pathway metabolites are compared to the factor's control (e.g. high-fat diet (HFD) compared to low-fat diet (LFD)). (e) Multiple factor signatures can be compared, such that distinct and shared signature pathways can be identified.

**Table S1.** Table of body mass gain from the beginning to the end of the diet intervention, the epididymal adipose depot mass at the end of the diet intervention, and blood glucose results from intraperitoneal glucose tolerance testing (ipGTT) without outliers presented as mean ± SD.

| | Female | | | | Male | | | | Main Effects | 2-way Interactions | 3-way Interaction | Main Effect F-values | 2-way Interactions F-values | 3-way Interaction F-values | Degrees of Freedom* | Error | Total |
|---|---|---|---|---|---|---|---|---|---|---|---|---|---|---|---|---|---|
| | Young (5-month) | | Aging (22-month) | | Young (5-month) | | Aging (22-month) | | | | | | | | | | |
| | LFD | HFD | LFD | HFD | LFD | HFD | LFD | HFD | | | | | | | | | |
| **Body Mass Gain from Baseline (%)** | (*n* = 9)<br>13.7 ± 3.9 | (*n* = 10)<br>13.2 ± 5.5 | (*n* = 8)<br>1.10 ± 3.66 | (*n* = 7)<br>7.82 ± 9.08 | (n = 10)<br>17.0 ± 7.9 | (n = 11)<br>53.0 ± 10.4 | (n = 8)<br>-9.89 ± 3.89 | (n = 8)<br>2.38 ± 4.81 | NS | NS | **Glucose AUC (p < 0.001)**<br>Sex*Age*Diet (p < 0.001) | NS | NS | **Glucose AUC (14.52)**<br>Sex*Age*Diet (15.60) | 8** | 61 | 69 |
| **Epididymal Adipose Depot Mass (g)** | (*n* = 10)<br>0.386 ± 0.156 | (*n* = 9)<br>0.491 ± 0.073 | (*n* = 9)<br>0.686 ± 0.269 | (*n* = 7)<br>0.787 ± 0.491 | (*n* = 10)<br>0.849 ± 0.258 | (*n* = 11)<br>2.08 ± 0.41 | (*n* = 8)<br>0.504 ± 0.121 | (*n* = 8)<br>0.909 ± 0.288 | NS | **Glucose AUC (p < 0.001)**<br>Sex*Age (p < 0.001)<br>Sex*Diet (p < 0.001)<br>Age*Diet (p = 0.037) | NS | NS | **Glucose AUC (26.11)**<br>Sex*Age (15.16)<br>Sex*Diet (15.04)<br>Age*Diet (4.57) | NS | 8** | 62 | 70 |
| **Glucose AUC** | (*n* = 9)<br>26000 ± 3320 | (*n* = 10)<br>28200 ± 3260 | (*n* = 9)<br>23600 ± 4650 | (*n* = 7)<br>30100 ± 7880 | (n = 10)<br>30800 ± 3520 | (n = 11)<br>44300 ± 4640 | (n = 8)<br>20700 ± 3340 | (n = 9)<br>28500 ± 9800 | NS | Sex*Age (p < 0.001)<br>Sex*Diet (p = 0.017)<br>Age*Diet (NS) | Sex*Age*Diet (NS, p = 0.055) | NS | Sex*Age (24.86)<br>Sex*Diet (6.06)<br>Age*Diet (NS) | Sex*Age*Diet (NS, 3.83) | 7 | 65 | 72 |
| **Baseline Glucose ($mg\ dL^{-1}$)** | (*n* = 10)<br>140 ± 25 | (*n* = 10)<br>152 ± 25 | (*n* = 9)<br>120 ± 15 | (*n* = 7)<br>149 ± 41 | (*n* = 10)<br>138 ±22 | (*n* = 11)<br>153 ± 24 | (*n* = 8)<br>104 ± 18 | (*n* = 9)<br>113 ± 25 | Age (NS)<br>Diet (p = 0.007)<br>Sex (NS) | Sex*Age (p = 0.024) | NS | Age (NS)<br>Diet (7.73)<br>Sex (NS) | Sex*Age (5.35) | NS | 7 | 66 | 73 |
| **Maximum Glucose ($mg\ dL^{-1}$)** | (*n* = 10)<br>384 ± 116 | (*n* = 10)<br>445 ± 58 | (*n* = 9)<br>329 ± 52 | (*n* = 7)<br>476 ± 58 | (n = 10)<br>369 ± 37 | (n = 11)<br>461 ± 13 | (n = 8)<br>303 ± 61 | (n = 9)<br>381 ± 121 | Age (p = 0.017)<br>Diet (p < 0.001)<br>Sex (NS, p = 0.089) | Sex*Age (NS, p = 0.088) | NS | Age (5.95)<br>Diet (29.07)<br>Sex (NS, 2.99) | Sex*Age (2.99) | NS | 7 | 66 | 73 |
| **Delta Glucose ($mg\ dL^{-1}$)** | (*n* = 10)<br>245 ± 123 | (*n* = 9)<br>294 ± 60 | (*n* = 9)<br>208 ± 44 | (*n* = 7)<br>327 ± 34 | (*n* = 10)<br>231 ± 53 | (*n* = 11)<br>307 ± 48 | (*n* = 8)<br>200 ± 57 | (*n* = 9)<br>268 ± 111 | Age (NS)<br>Diet (p < 0.001)<br>Sex (NS) | NS | NS | Age (NS)<br>Diet (20.28)<br>Sex (NS) | NS | NS | 7 | 66 | 73 |

**p- and F-values are reported for p < 0.10. Statistical significance is defined as p < 0.05. NS = No significance**

***Degrees of freedom for 3-way ANOVA (1 for each of 3 factors, 1 for each 2-way interaction between 3 factors, 1 for a 3-way interaction)**

****Degrees of freedom for 3-way ANCOVA (1 for a covariate, 1 for each of 3 factors, 1 for each 2-way interaction between 3 factors, 1 for a 3-way interaction)**

**LFD = Low-fat Diet HFD = High-fat Diet**

**Aged HFD Females had smaller n because 2 mice were excluded from all analyses due to tumors after diet intervention**
**Aged LFD Females had smaller n because 1 mouse died before the end of the diet intervention**
**Aged LFD Males had smaller n because 1 mouse died before the end of the diet intervention**

**Table S2.** Table of blood serum biomarker results without outliers presented as mean ± SD.

| | Female | | | | Male | | | | Main Effects | 2-way Interactions | 3-way Interaction | Main Effect F-values | 2-way Interactions F-values | 3-way Interaction F-values | Degrees of Freedom* | Error | Total |
|---|---|---|---|---|---|---|---|---|---|---|---|---|---|---|---|---|---|
| | Young (5-month) | | Aging (22-month) | | Young (5-month) | | Aging (22-month) | | | | | | | | | | |
| | LFD | HFD | LFD | HFD | LFD | HFD | LFD | HFD | | | | | | | | | |
| **IL-4 (ng/mL)** | (*n* = 10)<br>11.8 ± 7.7 | (*n* = 10)<br>14.9 ± 8.1 | (*n* = 9)<br>18.4 ± 8.8 | (*n* = 6)<br>15.1 ± 9.5 | (*n* = 10)<br>17.7 ± 8.8 | (*n* = 11)<br>19.5 ± 7.2 | (*n* = 8)<br>21.0 ± 7.7 | (*n* = 9)<br>17.0 ± 7.9 | Age (NS)<br>Diet (NS)<br>Sex (p = 0.033) | Age*Diet (NS, p = 0.088) | NS | Age (NS)<br>Diet (NS)<br>Sex (4.73) | Age*Diet (NS, 3.00) | NS | 7 | 65 | 72 |
| **IL-6 (pg/mL)** | (*n* = 9)<br>2.01 ± 0.31 | (*n* = 7)<br>1.93 ± 0.27 | (*n* = 9)<br>4.78 ± 3.15 | (*n* = 6)<br>3.72 ± 0.88 | (*n* = 7)<br>2.52 ± 1.68 | (*n* = 7)<br>1.81 ± 0.33 | (*n* = 6)<br>7.23 ± 3.63 | (*n* = 7)<br>3.36 ± 2.31 | Age (p < 0.001)<br>Diet (0.047)<br>Sex (NS) | Sex*Diet (NS, p = 0.062) | NS | Age (43.81)<br>Diet (4.15)<br>Sex (NS) | Sex*Diet (NS, 3.65) | NS | 7 | 50 | 57 |
| **IL-10 (pg/mL)** | (*n* = 10)<br>1.49 ± 0.39 | (*n* = 9)<br>1.38 ± 0.68 | (*n* = 9)<br>6.74 ± 6.89 | (*n* =7)<br>4.65 ± 3.21 | (*n* = 10)<br>1.10 ± 0.40 | (*n* = 9)<br>1.11 ± 0.41 | (*n* = 7)<br>2.47 ± 1.50 | (*n* = 8)<br>1.38 ± 0.69 | NS | Sex*Age (p = 0.036) | NS | NS | Sex*Age (4.60) | NS | 7 | 61 | 68 |
| **FGF21 (ng/mL)** | (*n* = 10)<br>582 ± 358 | (*n* = 10)<br>375 ± 180 | (*n* = 8)<br>754 ± 544 | (*n* = 7)<br>831 ± 532 | (*n* = 10)<br>972 ± 419 | (*n* = 11)<br>774 ± 332 | (*n* = 8)<br>1326 ± 968 | (*n* = 9)<br>737 ± 455 | Age (NS, p = 0.066)<br>Diet (NS)<br>Sex (p = 0.004) | Sex*Age (NS, p = 0.088) | NS | Age (NS, 3.49)<br>Diet (NS)<br>Sex (8.69) | Sex*Age (NS, 2.99) | NS | 7 | 65 | 72 |
| **Leptin (ng/mL)** | (*n* = 10)<br>70.8 ± 65.5 | (*n* = 9)<br>115 ± 45 | (*n* = 9)<br>119 ± 113 | (*n* = 6)<br>126 ± 105 | (*n* = 10)<br>251 ± 247 | (*n* = 11)<br>4340 ± 2600 | (*n* = 6)<br>18.2 ± 10.0 | (*n* = 8)<br>131 ± 140 | NS | NS | Sex*Age*Diet<br>(p = 0.018) | NS | NS | Sex*Age*Diet<br>(5.87) | 7 | 61 | 68 |
| **VEGF (pg/mL)** | (*n* = 10)<br>8.91 ± 2.52 | (*n* = 9)<br>7.58 ± 2.47 | (*n* = 9)<br>13.8 ± 3.42 | (*n* = 7)<br>15.5 ± 4.6 | (*n* = 10)<br>12.0 ± 1.3 | (*n* = 11)<br>10.6 ± 1.4 | (*n* = 8)<br>17.7 ± 7.04 | (*n* = 9)<br>11.6 ± 4.0 | NS | NS | Sex*Age*Diet<br>(p = 0.032) | NS | NS | Sex*Age*Diet<br>(4.79) | 7 | 65 | 72 |
| **Adiponectin (μg/mL)** | (*n* = 10)<br>11.5 ± 5.6 | (*n* = 9)<br>10.8 ± 2.4 | (*n* = 9)<br>11.6 ± 5.4 | (*n* = 7)<br>12.6 ± 4.4 | (*n* = 10)<br>7.18 ± 3.38 | (*n* = 10)<br>7.50 ± 2.95 | (*n* = 8)<br>10.9 ± 6.1 | (*n* = 9)<br>8.87 ± 3.29 | Age (NS, p = 0.098)<br>Diet (NS)<br>Sex (p = 0.005) | NS | NS | Age (NS, 2.83)<br>Diet (NS)<br>Sex (8.56) | NS | NS | 7 | 64 | 71 |
| **Resistin (ng/mL)** | (*n* = 10)<br>43.1 ± 15.8 | (*n* = 10)<br>36.3 ± 16.2 | (*n* = 9)<br>39.7 ± 12.3 | (*n* = 7)<br>42.4 ± 17.1 | (*n* = 10)<br>27.0 ± 5.2 | (*n* = 11)<br>27.4 ± 5.0 | (*n* = 8)<br>31.2 ± 4.9 | (*n* = 9)<br>19.9 ± 4.0 | NS | NS | Sex*Age*Diet<br>(p = 0.011) | NS | NS | Sex*Age*Diet<br>(6.78) | 7 | 66 | 73 |
| **Adipsin (μg/mL)** | (*n* = 10)<br>13.0 ± 2.7 | (*n* = 10)<br>15.3 ± 1.8 | (*n* = 9)<br>11.4 ± 2.3 | (*n* = 6)<br>13.0 ± 1.9 | (*n* = 9)<br>9.91 ± 1.25 | (*n* = 11)<br>6.84 ± 2.65 | (*n* = 7)<br>13.5 ± 7.0 | (*n* = 9)<br>9.91 ± 2.56 | NS | Sex*Age (p < 0.001)<br>Sex*Diet (p < 0.001) | NS | NS | Sex*Age (12.39)<br>Sex*Diet (13.14) | NS | 7 | 63 | 70 |
| **CRP (μg/mL)** | (*n* = 10)<br>4.09 ± 0.68 | (*n* = 10)<br>4.18 ± 0.47 | (*n* = 9)<br>4.70 ± 0.78 | (*n* = 7)<br>4.94 ± 0.74 | (*n* = 10)<br>5.19 ± 0.75 | (*n* = 11)<br>4.74 ± 0.92 | (*n* = 7)<br>3.65 ± 1.06 | (*n* = 9)<br>4.35 ± 1.07 | NS | Sex*Age (p < 0.001) | NS | NS | Sex*Age (17.89) | NS | 7 | 65 | 72 |

**p- and F-values are reported for p < 0.10. Statistical significance is defined as p < 0.05. NS = No significance**

***Degrees of freedom for 3-way ANOVA (1 for each of 3 factors, 1 for each 2-way interaction between 3 factors, 1 for a 3-way interaction)**

**LFD = Low-fat Diet** **HFD = High-fat Diet**

**Aged HFD Females had smaller n because 2 mice were excluded from all analyses due to tumors after diet intervention**
**Aged LFD Females had smaller n because 1 mouse died before the end of the diet intervention**
**Aged LFD Males had smaller n because 1 mouse died before the end of the diet intervention**

**Table S3.** Micro-computed tomography results for the microarchitecture of the left femora without outliers presented as mean ± SD.

| | Female | | | | Male | | | | Main Effects | 2-way Interactions | 3-way Interaction | Main Effect F-values | 2-way Interactions F-values | 3-way Interaction F-values | Degrees of Freedom* | Error | Total |
|---|---|---|---|---|---|---|---|---|---|---|---|---|---|---|---|---|---|
| | Young (5-month) | | Aging (22-month) | | Young (5-month) | | Aging (22-month) | | | | | | | | | | |
| | LFD | HFD | LFD | HFD | LFD | HFD | LFD | HFD | | | | | | | | | |
| **BV/TV (%)** | (*n* = 10)<br>6.31 ± 2.15 | (*n* = 9)<br>4.74 ± 1.03 | (*n* = 9)<br>2.07 ± 1.53 | (*n* = 6)<br>1.86 ± 0.72 | (*n* = 10)<br>14.3 ± 5.1 | (*n* = 10)<br>14.4 ± 1.9 | (*n* = 8)<br>7.78 ± 1.70 | (*n* = 9)<br>9.98 ± 4.51 | Age (p < 0.001)<br>Diet (NS)<br>Sex (p < 0.001) | NS | NS | Age (62.14)<br>Diet (NS)<br>Sex (174.95) | NS | NS | 7 | 63 | 70 |
| **BMD (mgHA/$cm^3$)** | (*n* = 10)<br>117± 20 | (*n* = 9)<br>96.9 ± 12.3 | (*n* = 9)<br>69.6 ± 12.2 | (*n* = 6)<br>60.0 ± 14.0 | (*n* = 10)<br>184± 42 | (*n* = 10)<br>188 ± 15 | (*n* = 8)<br>124 ± 13 | (*n* = 9)<br>141 ± 41 | Age (p < 0.001) | Sex*Age (NS, p = 0.081)<br>Sex*Diet (p = 0.010) | NS | Age (86.43) | Sex*Age (NS, 3.14)<br>Sex*Diet (7.04) | NS | 7 | 63 | 70 |
| **Conn. D. (1/$mm^3$)** | (*n* = 10)<br>25.5 ± 14.2 | (*n* = 9)<br>14.7 ± 6.5 | (*n* = 8)<br>1.70 ± 1.81 | (*n* = 6)<br>1.70 ± 1.72 | (*n* = 10)<br>94.0 ± 54.5 | (*n* = 10)<br>110 ± 16 | (*n* = 8)<br>30.0 ± 17.3 | (*n* = 9)<br>47.6 ± 32.8 | Age (p < 0.001)<br>Diet (NS)<br>Sex (p < 0.001) | Sex*Diet (NS, p = 0.074) | NS | Age (88.70)<br>Diet (NS)<br>Sex (170.59) | Sex*Diet (NS, 3.30) | NS | 7 | 62 | 69 |
| **Tb. N. (1/mm)** | (*n* = 9)<br>3.06 ± 0.18 | (*n* = 10)<br>2.75 ± 0.29 | (*n* = 8)<br>1.33 ± 0.16 | (*n* = 6)<br>1.44 ± 0.22 | (*n* = 10)<br>4.12 ± 1.00 | (*n* = 10)<br>4.66 ± 0.23 | (*n* = 8)<br>2.86 ± 0.49 | (*n* = 9)<br>3.13 ± 0.86 | NS | Sex*Age (p < 0.001) | NS | NS | Sex*Age (18.54) | NS | 7 | 62 | 69 |
| **Tb. Th. (mm)** | (*n* = 10)<br>0.054 ± 0.008 | (*n* = 10)<br>0.054 ± 0.004 | (*n* = 9)<br>0.059 ± 0.008 | (*n* = 6)<br>0.065 ± 0.010 | (*n* = 9)<br>0.054 ± 0.002 | (*n* = 11)<br>0.052 ± 0.003 | (*n* = 8)<br>0.059 ± 0.007 | (*n* = 9)<br>0.056 ± 0.004 | Age (p < 0.001) | Sex*Diet (p = 0.048) | NS | Age (18.74) | Sex*Diet (4.06) | NS | 7 | 64 | 71 |
| **Tb. Sp (mm)** | (*n* = 9)<br>0.328 ± 0.022 | (*n* = 10)<br>0.368 ± 0.038 | (*n* = 9)<br>0.737 ± 0.131 | (*n* = 6)<br>0.708 ± 0.116 | (*n* = 10)<br>0.251 ± 0.075 | (*n* = 10)<br>0.206 ± 0.012 | (*n* = 8)<br>0.350 ± 0.057 | (*n* = 9)<br>0.334 ± 0.094 | NS | Sex*Age (p = 0.001)<br>Sex*Diet (NS, p = 0.084) | NS | NS | Sex*Age (12.97)<br>Sex*Diet (NS, 3.08) | NS | 7 | 63 | 70 |
| **Ct. Ar ($mm^2$)** | (*n* = 10)<br>0.838 ± 0.043 | (*n* = 9)<br>0.808 ± 0.028 | (*n* = 9)<br>0.834 ± 0.074 | (*n* = 7)<br>0.828 ± 0.069 | (*n* = 10)<br>0.842 ± 0.055 | (*n* = 11)<br>0.800 ± 0.036 | (*n* = 8)<br>0.908 ± 0.068 | (*n* = 9)<br>0.876 ± 0.082 | Diet (NS, p = 0.052) | Sex*Age (p = 0.028) | NS | Diet (NS, 3.93) | Sex*Age (5.04) | NS | 7 | 65 | 72 |
| **Ct. Th (mm)** | (*n* = 9)<br>0.206 ± 0.005 | (*n* = 9)<br>0.195 ± 0.005 | (*n* = 8)<br>0.176 ± 0.009 | (*n* = 7)<br>0.170 ± 0.021 | (*n* = 10)<br>0.182 ± 0.012 | (*n* = 11)<br>0.182 ± 0.008 | (*n* = 8)<br>0.166 ± 0.014 | (*n* = 9)<br>0.162 ± 0.018 | Age (p < 0.001)<br>Diet (NS, 0.072)<br>Sex (p < 0.001) | NS | NS | Age (63.76)<br>Diet (NS, 3.35)<br>Sex (22.78) | NS | NS | 7 | 63 | 70 |
| **Ct. TMD (mgHA/$cm^3$)** | (*n* = 10)<br>1239 ± 12 | (*n* = 10)<br>1235± 11 | (*n* = 9)<br>1238 ± 16 | (*n* = 7)<br>1237 ± 20 | (*n* = 10)<br>1212 ± 43 | (*n* = 11)<br>1220 ± 13 | (*n* = 8)<br>1219 ± 18 | (*n* = 9)<br>1212 ± 17 | Age (NS)<br>Diet (NS)<br>Sex (p < 0.001) | NS | NS | Age (NS)<br>Diet (NS)<br>Sex (18.89) | NS | NS | 7 | 66 | 73 |
| **Ct. Porosity (%)** | (*n* = 9)<br>0.649 ± 0.076 | (*n* = 9)<br>0.625 ± 0.054 | (*n* = 8)<br>0.717 ± 0.143 | (*n* = 5)<br>0.677 ± 0.052 | (*n* = 9)<br>0.853 ± 0.185 | (*n* = 11)<br>1.09 ± 0.29 | (*n* = 7)<br>0.922 ± 0.281 | (*n* = 8)<br>0.835 ± 0.119 | NS | Sex*Age (p = 0.035) | NS | NS | Sex*Age (4.68) | NS | 7 | 58 | 65 |
| **Ma. Ar ($mm^2$)** | (*n* = 10)<br>0.814 ± 0.061 | (*n* = 9)<br>0.847 ± 0.054 | (*n* = 9)<br>1.34 ± 0.12 | (*n* = 7)<br>1.35 ± 0.17 | (*n* = 10)<br>1.07 ± 0.21 | (*n* = 11)<br>0.934 ± 0.072 | (*n* = 8)<br>1.68 ± 0.19 | (*n* = 9)<br>1.68 ± 0.21 | NS | Sex*Age (p = 0.023) | NS | NS | Sex*Age (5.40) | NS | 7 | 65 | 72 |
| **pMOI ($mm^4$)** | (*n* = 10)<br>0.345 ± 0.039 | (*n* = 10)<br>0.336 ± 0.023 | (*n* = 9)<br>0.476 ± 0.057 | (*n* = 7)<br>0.471 ± 0.060 | (*n* = 9)<br>0.410 ± 0.060 | (*n* = 11)<br>0.363 ± 0.033 | (*n* = 8)<br>0.653 ± 0.085 | (*n* = 9)<br>0.623 ± 0.084 | Diet (NS, p = 0.098) | Sex*Age (p < 0.001) | NS | Diet (NS, 2.82) | Sex*Age (19.45) | NS | 7 | 65 | 72 |
| **Imin ($mm^4$)** | (*n* = 10)<br>0.118 ± 0.010 | (*n* = 10)<br>0.118 ± 0.010 | (*n* = 9)<br>0.184 ± 0.022 | (*n* = 7)<br>0.186 ± 0.027 | (*n* = 9)<br>0.127 ± 0.015 | (*n* = 11)<br>0.114 ± 0.010 | (*n* = 8)<br>0.207 ± 0.034 | (*n* = 9)<br>0.202 ± 0.025 | Age (p < 0.001)<br>Diet (NS)<br>Sex (NS, p = 0.072) | NS | NS | Age (312.77)<br>Diet (NS)<br>Sex (NS, 3.34) | NS | NS | 7 | 65 | 72 |

**p- and F-values are reported for p < 0.10. Statistical significance is defined as p < 0.05. NS = No significance**

***Degrees of freedom for 3-way ANOVA (1 for each of 3 factors, 1 for each 2-way interaction between 3 factors, 1 for a 3-way interaction)**

**LFD = Low-fat Diet HFD = High-fat Diet**

**Aged HFD Females had smaller n because 2 mice were excluded from all analyses due to tumors after diet intervention**
**Aged LFD Females had smaller n because 1 mouse died before the end of the diet intervention**
**Aged LFD Males had smaller n because 1 mouse died before the end of the diet intervention**

**Table S4.** Biomechanical results from notched fracture testing (right femora) and three-point bending (left femora) without outliers. Presented as mean ± SD.

| | Female | | | | Male | | | | Main Effects | 2-way Interactions | 3-way Interaction | Main Effect F-values | 2-way Interactions F-values | 3-way Interaction F-values | Degrees of Freedom* | Error | Total |
|---|---|---|---|---|---|---|---|---|---|---|---|---|---|---|---|---|---|
| | Young (5-month) | | Aging (22-month) | | Young (5-month) | | Aging (22-month) | | | | | | | | | | |
| | LFD | HFD | LFD | HFD | LFD | HFD | LFD | HFD | | | | | | | | | |
| **Notched Fracture Toughness Testing** | | | | | | | | | | | | | | | | | |
| **$Kc_{max}$ (mPa·m$^{1/2}$)** | (*n = 8*)<br>4.83 ± 0.53 | (*n = 7*)<br>4.19 ± 0.53 | (*n = 8*)<br>3.57 ± 0.78 | (*n = 5*)<br>3.36 ± 0.61 | (*n = 9*)<br>4.27 ± 0.53 | (*n = 7*)<br>3.83 ± 1.22 | (*n = 5*)<br>4.06 ± 0.68 | (*n = 7*)<br>3.83 ± 0.57 | Diet (NS, p = 0.056) | Sex*Age (p = 0.019) | NS | Diet (NS, 3.84) | Sex*Age (5.86) | NS | 7 | 48 | 55 |
| **$Kc_{init}$ (mPa·m$^{1/2}$)** | (*n = 8*)<br>3.54 ± 0.53 | (*n = 7*)<br>3.24 ± 0.65 | (*n = 8*)<br>2.68 ± 0.94 | (*n = 5*)<br>2.32 ± 0.52 | (*n = 9*)<br>3.71 ± 0.65 | (*n = 7*)<br>3.14 ± 1.29 | (*n = 5*)<br>3.65 ± 0.69 | (*n = 7*)<br>2.82 ± 0.82 | Age (p = 0.017)<br>Diet (p = 0.023)<br>Sex (NS, p = 0.085) | NS | NS | Age (6.12)<br>Diet (5.49)<br>Sex (NS, 3.09) | NS | NS | 7 | 48 | 55 |
| **Three-Point Bending** | | | | | | | | | | | | | | | | | |
| **Elastic Modulus (GPa)** | (*n = 9*)<br>7.77 ± 1.47 | (*n = 10*)<br>8.08 ± 1.57 | (*n = 9*)<br>5.63 ± 0.99 | (*n = 7*)<br>5.12 ± 1.16 | (*n = 10*)<br>6.24 ± 0.97 | (*n = 11*)<br>6.17 ± 0.94 | (*n = 8*)<br>4.08 ± 1.27 | (*n = 8*)<br>3.82 ± 0.74 | Age (p < 0.001)<br>Diet (NS)<br>Sex (p < 0.001) | NS | NS | Age (74.13)<br>Diet (NS)<br>Sex (31.78) | NS | NS | 7 | 64 | 71 |
| **Ultimate Strength (MPa)** | (*n = 9*)<br>192 ± 21 | (*n = 10*)<br>207± 16 | (*n = 9*)<br>131 ± 20 | (*n = 7*)<br>146 ± 21 | (*n = 9*)<br>182 ± 17 | (*n = 11*)<br>172 ± 18 | (*n = 8*)<br>130 ± 32 | (*n = 8*)<br>121 ± 26 | Age (p < 0.001) | Sex*Diet (p = 0.021) | NS | Age (121.36) | Sex*Diet (5.58) | NS | 7 | 63 | 70 |
| **Yield Strength (MPa)** | (*n = 9*)<br>172 ± 27 | (*n = 10*)<br>157 ± 26 | (*n = 9*)<br>82.5 ± 13.3 | (*n = 7*)<br>101 ± 33 | (*n = 9*)<br>155 ± 39 | (*n = 11*)<br>145 ± 24 | (*n = 7*)<br>93.2 ± 24.8 | (*n =8*)<br>72.5 ± 15.2 | Age (p < 0.001)<br>Diet (NS)<br>Sex (NS) | NS | Sex*Age*Diet<br>(NS, p = 0.091) | Age (118.97)<br>Diet (NS)<br>Sex (NS) | NS | Sex*Age*Diet<br>(NS, 2.94) | 7 | 62 | 69 |
| **Stiffness (N /mm)** | (*n = 8*)<br>92.7 ± 8.8 | (*n = 10*)<br>89.9 ± 19.4 | (*n = 9*)<br>96.5 ± 18.0 | (*n = 7*)<br>87.6 ± 15.2 | (*n = 10*)<br>77.0 ± 12.2 | (*n = 11*)<br>65.9 ± 11.7 | (*n = 8*)<br>78.0 ± 24.6 | (*n = 8*)<br>72.1 ± 12.9 | Age (NS)<br>Diet (NS, p = 0.065)<br>Sex (p = 0.001) | NS | NS | Age (NS)<br>Diet (NS, 3.52)<br>Sex (23.24) | NS | NS | 7 | 63 | 70 |
| **Post Yield Strain** | (*n = 9*)<br>0.034 ± 0.039 | (*n = 9*)<br>0.031 ± 0.015 | (*n = 9*)<br>0.040 ± 0.038 | (*n = 7*)<br>0.023 ± 0.014 | (*n = 9*)<br>0.017 ± 0.012 | (*n = 11*)<br>0.039 ± 0.029 | (*n = 8*)<br>0.033 ± 0.042 | (*n = 8*)<br>0.098 ± 0.099 | Age (NS)<br>Diet (p = 0.036)<br>Sex (NS) | Sex*Diet (NS, p = 0.091) | NS | Age (NS)<br>Diet (4.57)<br>Sex (NS) | Sex*Diet (NS, 2.95) | NS | 7 | 62 | 69 |
| **Energy-at-Fracture (mJ)** | (*n = 9*)<br>5.43 ± 2.99 | (*n = 10*)<br>6.00 ± 2.99 | (*n = 9*)<br>3.85 ± 2.94 | (*n = 7*)<br>3.14 ± 1.56 | (*n = 10*)<br>4.01 ± 1.28 | (*n = 11*)<br>4.06 ± 1.29 | (*n = 8*)<br>4.62 ± 3.99 | (*n = 8*)<br>4.80 ± 3.12 | Age (p = 0.032)<br>Diet (NS)<br>Sex (NS) | Sex*Age (NS, p = 0.059) | NS | Age (4.83)<br>Diet (NS)<br>Sex (NS) | Sex*Age (NS, 3.69) | NS | 7 | 64 | 71 |

**p- and F-values are reported for p < 0.10. Statistical significance is defined as p < 0.05. NS = No significance**
***Degrees of freedom for 3-way ANOVA (1 for each of 3 factors, 1 for each 2-way interaction between 3 factors, 1 for a 3-way interaction)**

**LFD = Low-fat Diet HFD = High-fat Diet**

**Aged HFD Females had smaller total n because 2 mice were excluded from all analyses due to tumors after diet intervention**
**Aged LFD Females had smaller total n because 1 mouse died before the end of the diet intervention**
**Aged LFD Males had smaller total n because 1 mouse died before the end of the diet intervention**

**Fracture Toughness Testing - Groups had smaller n due disqualification of samples that had errors in notch angles compared to loading axis and movement during testing**
**Three-point Bending - Groups had smaller n due to disqualification of samples that moved during testing**

**Table S5.** Bone turnover results from quantitative histomorphometry of proximal left femora. Presented as mean ± SD.

| | Female | | | | Male | | | | Main Effects | 2-way Interactions | 3-way Interaction | Main Effect F-values | 2-way Interactions F-values | 3-way Interaction F-values | Degrees of Freedom* | Error | Total |
|---|---|---|---|---|---|---|---|---|---|---|---|---|---|---|---|---|---|
| | Young (5-month) | | Aging (22-month) | | Young (5-month) | | Aging (22-month) | | | | | | | | | | |
| | LFD | HFD | LFD | HFD | LFD | HFD | LFD | HFD | | | | | | | | | |
| | *(n = 5)* | *(n = 9)* | *(n = 9)* | *(n = 6)* | *(n = 9)* | *(n = 11)* | *(n = 6)* | *(n = 8)* | | | | | | | | | |
| **MS / BS (%)** | 40.1 ± 13.8 | 37.8 ± 6.2 | 17.0 ± 8.5 | 16.9 ± 8.1 | 21.8 ± 7.2 | 24.4 ± 6.0 | 9.49 ± 4.96 | 11.4 ± 11.0 | NS | Sex*Age (p= 0.034) | NS | NS | Sex*Age (4.73) | NS | 7 | 55 | 62 |
| | (*n = 5*) | (*n = 9*) | (*n = 9*) | (*n = 6*) | (*n = 9*) | (*n = 11*) | (*n = 6*) | (*n = 8*) | | | | | | | | | |
| **E. MS / E. BS (%)** | 45.0 ± 20.8 | 28.8 ± 10.0 | 27.5 ± 19.2 | 25.9 ± 10.8 | 15.3 ± 9.8 | 6.39 ± 4.59 | 12.8 ± 7.4 | 9.12 ± 9.08 | Age (NS)<br>Diet (p = 0.019)<br>Sex (p < 0.001) | NS | NS | Age (NS)<br>Diet (5.82)<br>Sex (46.94) | NS | NS | 7 | 55 | 62 |
| | *(n = 5)* | *(n = 9)* | *(n = 9)* | *(n = 6)* | *(n = 9)* | *(n = 11)* | (*n = 7*) | (*n = 8*) | | | | | | | | | |
| **P. MS / P. BS(%)** | 37.4 ± 17.4 | 43.3 ± 11.9 | 9.42 ± 8.67 | 10.2 ± 8.0 | 25.7 ± 12.8 | 34.9 ± 9.15 | 8.67 ± 4.94 | 13.0 ± 12.5 | Age (p < 0.001)<br>Diet (NS, p = 0.076)<br>Sex (NS) | Sex*Age (NS, p = 0.053) | NS | Age (79.56)<br>Diet (NS, 3.26)<br>Sex (NS) | Sex*Age (NS, 3.91) | NS | 7 | 56 | 63 |

**p- and F-values are reported for p < 0.10. Statistical significance is defined as p < 0.05. NS = No significance**

***Degrees of freedom for 3-way ANOVA (1 for each of 3 factors, 1 for each 2-way interaction between 3 factors, 1 for a 3-way interaction)**

**LFD = Low-fat Diet** **HFD = High-fat Diet**

**Young LFD Females have smaller n because 5 proximal femurs after three-point bending were not sufficient for accurate measurement of labels**
**Aged HFD Females had smaller n because 2 mice were excluded from all analyses due to tumors after diet intervention**
**Aged LFD Females had smaller n because 1 mouse died before the end of the diet intervention**
**Aged LFD Males had smaller n because 1 mouse died before the end of the diet intervention**

**Table S6.** Results related to bone turnover. *Opg* and *Rankl* are from qRT-PCR (marrow-flushed left tibiae) and blood P1NP and CTX1 are from blood serum biomarkers. Presented as mean ± SD.

| | Female | | | | Male | | | | Main Effects | 2-way Interactions | 3-way Interaction | Main Effect F-values | 2-way Interactions F-values | 3-way Interaction F-values | Degrees of Freedom* | Error | Total |
|---|---|---|---|---|---|---|---|---|---|---|---|---|---|---|---|---|---|
| | Young (5-month) | | Aging (22-month) | | Young (5-month) | | Aging (22-month) | | | | | | | | | | |
| | LFD | HFD | LFD | HFD | LFD | HFD | LFD | HFD | | | | | | | | | |
| ***Opg:Rankl* Ratio** | (*n* = 7) 1.94 ± 0.99 | (*n* = 10) 4.56 ± 4.33 | (*n* = 9) 4.02 ± 3.90 | (*n* = 7) 1.10 ± 0.67 | (*n* = 9) 4.72 ± 3.92 | (*n* = 8) 3.25 ± 1.45 | (*n* = 6) 2.15 ± 1.23 | (*n* = 8) 1.12 ± 0.64 | NS | Age*Diet (p = 0.013) | NS | NS | Age*Diet (6.56) | NS | 7 | 56 | 63 |
| ***Opg*** | (*n* = 9) 1.18 ± 0.75 | (*n* = 9) 1.14 ± 1.37 | (*n* = 8) 0.75 ± 0.85 | (*n* = 7) 0.62 ± 0.53 | (*n* = 9) 0.84 ± 0.70 | (*n* = 7) 0.61 ± 0.14 | (*n* = 6) 0.74 ± 0.65 | (*n* = 8) 0.36 ± 0.37 | Age (p = 0.010)<br>Diet (NS)<br>Sex (NS) | NS | NS | Age (7.07)<br>Diet (NS)<br>Sex (NS) | NS | NS | 7 | 55 | 62 |
| ***Rankl*** | (*n* = 9) 1.24 ± 0.81 | (*n* = 10) 0.98 ± 0.70 | (*n* = 8) 0.59 ± 0.44 | (*n* = 7) 1.92 ± 1.81 | (*n* = 8) 0.44 ± 0.24 | (*n* = 8) 0.59 ± 0.31 | (*n* = 6) 0.87 ± 0.53 | (*n* = 8) 2.27 ± 3.38 | NS | NS | NS | NS | NS | NS | 7 | 56 | 63 |
| **P1NP:CTX1 Ratio** | (*n* = 9) 16.3 ± 7.5 | (*n* = 10) 32.2 ± 23.2 | (*n* = 9) 58.6 ± 78.0 | (*n* = 6) 27.0 ± 10.9 | (*n* = 10) 24.8 ± 15.8 | (*n* = 11) 32.0 ± 20.4 | (*n* = 8) 21.2 ± 16.6 | (*n* = 8) 19.2 ± 7.4 | NS | NS | NS | NS | NS | NS | 7 | 63 | 70 |
| **P1NP (ng/mL)** | (*n* = 10) 53.1 ± 27.0 | (*n* = 10) 67.7 ± 55.8 | (*n* = 8) 57.7 ± 26.1 | (*n* = 6) 78.8 ± 28.8 | (*n* = 10) 56.4 ± 30.4 | (*n* = 11) 71.6 ± 39.7 | (*n* = 8) 42.7 ± 22.9 | (*n* = 9) 59.9 ± 29.3 | **Glucose AUC (p = 0.024)**<br>Age (NS)<br>Diet (NS)<br>Sex (NS) | NS | NS | **Glucose AUC (5.38)**<br>Age (NS)<br>Diet (NS)<br>Sex (NS) | NS | NS | 8** | 62 | 70 |
| **CTX1 (ng/mL)** | (*n* = 10) 2.92 ± 0.77 | (*n* = 10) 2.28 ± 0.88 | (*n* = 9) 2.44 ± 0.99 | (*n* = 7) 2.91 ± 0.90 | (*n* = 10) 2.58 ± 0.74 | (*n* = 10) 2.37 ± 0.45 | (*n* = 8) 2.38 ± 0.75 | (*n* = 8) 2.47 ± 0.53 | NS | Age*Diet (NS, p = 0.058) | NS | NS | Age*Diet (NS, 3.74) | NS | 7 | 64 | 71 |

**p- and F-values are reported for p < 0.10. Statistical significance is defined as p < 0.05. NS = No significance. Bolded Glucose AUC indicates a significant covariate with adjusted p-values.**
***Degrees of freedom for 3-way ANOVA (1 for each of 3 factors, 1 for each 2-way interaction between 3 factors, 1 for a 3-way interaction)**
****Degrees of freedom for 3-way ANCOVA (1 for a covariate, 1 for each of 3 factors, 1 for each 2-way interaction between 3 factors, 1 for a 3-way interaction)**

**LFD = Low-fat Diet HFD = High-fat Diet**

**Aged HFD Females had smaller n because 2 mice were excluded from all analyses due to tumors after diet intervention**
**Aged LFD Females had smaller n because 1 mouse died before the end of the diet intervention**
**Aged LFD Males had smaller n because 1 mouse died before the end of the diet intervention**

**Table S7.** Results for histology – TRAP-positive osteoclast and adipocyte number density and morphology from right tibiae. Presented as mean ± SD.

| | Female | | | | Male | | | | Main Effects | 2-way Interactions | 3-way Interaction | Main Effect F-values | 2-way Interactions F-values | 3-way Interaction F-values | Degrees of Freedom* | Error | Total |
|---|---|---|---|---|---|---|---|---|---|---|---|---|---|---|---|---|---|
| | Young (5-month) | | Aging (22-month) | | Young (5-month) | | Aging (22-month) | | | | | | | | | | |
| | LFD | HFD | LFD | HFD | LFD | HFD | LFD | HFD | | | | | | | | | |
| **Osteoclast Number Density (cells/mm)** | (*n* = 10) 5.91 ± 2.67 | (*n* = 10) 5.53 ± 3.02 | (*n* = 9) 6.00 ± 3.45 | (*n* = 7) 2.70 ± 2.80 | (*n* = 10) 0.364 ± 0.538 | (*n* = 11) 1.38 ± 0.46 | (*n* = 8) 0.804 ± 0.884 | (*n* = 7) 0.00 ± 0.00 | NS | Age*Diet (p < 0.001)<br>Sex*Diet (p = 0.036) | NS | NS | Age*Diet (15.79)<br>Sex*Diet (4.60) | NS | 7 | 64 | 71 |
| **Adipocyte Number Density (cells/mm$^2$)** | (*n* = 9) 121 ± 29 | (*n* = 9) 142 ± 45 | (*n* = 7) 250 ± 218 | (*n* = 6) 85.1 ± 52.8 | (*n* = 9) 69.3 ± 41.2 | (*n* = 10) 102 ± 48 | (*n* = 7) 30.4 ± 14.2 | (*n* = 5) 33.3 ± 33.0 | NS | Age*Diet (p = 0.021)<br>Sex*Age (p = 0.006) | NS | NS | Age*Diet (5.65)<br>Sex*Age (8.16) | NS | 7 | 54 | 61 |
| **Adipocyte size (mm$^2$)** | (*n* = 9) 4.51E-4 ± 1.48E-4 | (*n* = 10) 4.63E-4 ± 1.17E-4 | (*n* = 7) 5.04E-4 ± 1.16E-4 | (*n* = 7) 5.22E-4 ± 1.14E-4 | (*n* = 9) 6.73E-4 ± 2.83E-4 | (*n* = 11) 9.73E-4 ± 1.92E-4 | (*n* = 7) 4.79E-4 ± 1.37E-4 | (*n* = 6) 5.92E-4 ± 0.93E-4 | NS | Sex*Age (p < 0.001)<br>Sex*Diet (p = 0.026) | NS | NS | Sex*Age (16.74)<br>Sex*Diet (5.20) | NS | 7 | 58 | 65 |

**p- and F-values are reported for p < 0.10. Statistical significance is defined as p < 0.05. NS = No significance**
***Degrees of freedom for 3-way ANOVA (1 for each of 3 factors, 1 for each 2-way interaction between 3 factors, 1 for a 3-way interaction)**

**LFD = Low-fat Diet** **HFD = High-fat Diet**

**Aged HFD Females had smaller n because 2 mice were excluded from all analyses due to tumors after diet intervention**
**Aged LFD Females had smaller n because 1 mouse died before the end of the diet intervention**
**Aged LFD Males had smaller n because 1 mouse died before the end of the diet intervention**

**Table S8.** Osteocyte number density, TUNEL-positive osteocytes from histology of right tibiae. Presented as mean ± SD.

| | Female | | | | Male | | | | Main Effects | 2-way Interactions | 3-way Interaction | Main Effect F-values | 2-way Interactions F-values | 3-way Interaction F-values | Degrees of Freedom* | Error | Total |
|---|---|---|---|---|---|---|---|---|---|---|---|---|---|---|---|---|---|
| | Young (5-month) | | Aging (22-month) | | Young (5-month) | | Aging (22-month) | | | | | | | | | | |
| | LFD | HFD | LFD | HFD | LFD | HFD | LFD | HFD | | | | | | | | | |
| **TUNEL-positive Osteocytes (% filled lacunae)** | (*n = 10*) 7.18 ± 3.30 | (*n = 9*) 6.59 ± 2.66 | (*n = 8*) 5.78 ± 4.90 | (*n = 7*) 9.91 ± 5.12 | (*n = 10*) 6.00 ± 3.82 | (*n = 11*) 6.75 ± 3.14 | (*n = 8*) 1.50 ± 4.23 | (*n = 8*) 11.8 ± 5.9 | NS | Age*Diet (p = 0.024) | NS | NS | Age*Diet (5.33) | NS | 7 | 63 | 70 |
| **Osteocyte Number Density (cells/mm$^2$)** | (*n = 10*) 1380 ± 383 | (*n = 9*) 1300 ± 438 | (*n = 8*) 1370 ± 522 | (*n = 7*) 1440 ± 392 | (*n = 10*) 1400 ± 567 | (*n = 11*) 1800 ± 410 | (*n = 7*) 792 ± 196 | (*n = 8*) 1270 ± 412 | NS | Sex*Age (p = 0.001)<br>Sex*Diet (p = 0.014) | NS | NS | Sex*Age (11.59)<br>Sex*Diet (6.46) | NS | 7 | 62 | 69 |

**p- and F-values are reported for p < 0.10. Statistical significance is defined as p < 0.05. NS = No significance**

***Degrees of freedom for 3-way ANOVA (1 for each of 3 factors, 1 for each 2-way interaction between 3 factors, 1 for a 3-way interaction)**

**LFD = Low-fat Diet** **HFD = High-fat Diet**

**Aged HFD Females had smaller n because 2 mice were excluded from all analyses due to tumors after diet intervention**
**Aged LFD Females had smaller n because 1 mouse died before the end of the diet intervention**
**Aged LFD Males had smaller n because 1 mouse died before the end of the diet intervention**

**Table S9.** Lacunar-canicular system resorption markers from qRT-PCR on marrow-flushed left tibiae. Presented as mean ± SD.

| | Female | | | | Male | | | | Main Effects | 2-way Interactions | 3-way Interaction | Main Effect F-values | 2-way Interactions F-values | 3-way Interaction F-values | Degrees of Freedom* | Error | Total |
|---|---|---|---|---|---|---|---|---|---|---|---|---|---|---|---|---|---|
| | Young (5-month) | | Aging (22-month) | | Young (5-month) | | Aging (22-month) | | | | | | | | | | |
| | LFD | HFD | LFD | HFD | LFD | HFD | LFD | HFD | | | | | | | | | |
| ***Acp5*** | (*n* = 9)<br>1.18 ± 0.64 | (*n* = 10)<br>1.18 ± 0.93 | (*n* = 8)<br>1.34 ± 0.55 | (*n* = 6)<br>1.58 ± 1.29 | (*n* = 9)<br>0.69 ± 0.33 | (*n* = 8)<br>0.45 ± 0.23 | (*n* = 8)<br>1.12 ± 0.36 | (*n* = 5)<br>1.03 ± 0.85 | Age (p = 0.015)<br>Diet (NS)<br>Sex (p = 0.036) | NS | NS | Age (4.64)<br>Diet (NS)<br>Sex (6.36) | NS | NS | 7 | 55 | 62 |
| ***Mmp2*** | (*n* = 9)<br>1.34 ± 1.02 | (*n* = 10)<br>1.72 ± 1.39 | (*n* = 8)<br>0.49 ± 0.25 | (*n* = 6)<br>0.45 ± 0.32 | (*n* = 9)<br>0.80 ± 0.50 | (*n* = 8)<br>0.49 ± 0.26 | (*n* = 6)<br>0.85 ± 0.96 | (*n* = 8)<br>0.50 ± 0.55 | Age (p = 0.003)<br>Diet (NS)<br>Sex (NS) | NS | NS | Age (9.70)<br>Diet (NS)<br>Sex (NS) | NS | NS | 7 | 56 | 63 |
| ***Mmp13*** | (*n* = 9)<br>1.13 ± 0.55 | (*n* = 10)<br>1.21 ± 0.66 | (*n* = 8)<br>0.90 ± 0.41 | (*n* = 6)<br>1.45 ± 1.26 | (*n* = 9)<br>0.77 ± 0.30 | (*n* = 8)<br>0.44 ± 0.29 | (*n* = 6)<br>1.41 ± 1.02 | (*n* = 8)<br>0.94 ± 0.50 | NS | Sex*Age (NS, 0.055)<br>Sex*Diet (p = 0.047) | NS | NS | Sex*Age (NS, 3.85)<br>Sex*Diet (4.11) | NS | 7 | 56 | 63 |
| ***Mmp14*** | (*n* = 8)<br>1.17 ± 0.71 | (*n* = 7)<br>0.90 ± 0.49 | (*n* = 9)<br>0.78 ± 0.47 | (*n* = 9)<br>1.18 ± 1.15 | (*n* = 6)<br>0.76 ± 0.37 | (*n* = 8)<br>0.64 ± 0.22 | (*n* = 9)<br>1.01 ± 0.74 | (*n* = 9)<br>0.63 ± 0.35 | **Glucose AUC (p = 0.040)**<br>Age (NS)<br>Diet (NS)<br>Sex (NS) | NS | NS | **Glucose AUC (4.46)**<br>Age (NS)<br>Diet (NS)<br>Sex (NS) | NS | NS | 8** | 53 | 61 |
| ***Ctsk*** | (*n* = 8)<br>1.16 ± 0.61 | (*n* = 10)<br>2.38 ± 2.84 | (*n* = 8)<br>1.67 ± 1.08 | (*n* = 5)<br>2.36 ± 3.69 | (*n* = 9)<br>1.09 ± 0.85 | (*n* = 8)<br>0.73 ± 0.53 | (*n* = 6)<br>3.14 ± 2.15 | (*n* = 8)<br>2.05 ± 2.44 | NS | NS | NS | NS | NS | NS | 7 | 54 | 61 |
| ***Atp6v0d2*** | (*n* = 9)<br>1.08 ± 0.40 | (*n* = 9)<br>0.80 ± 0.29 | (*n* = 8)<br>0.58 ± 0.35 | (*n* = 7)<br>1.65 ± 1.35 | (*n* = 9)<br>0.87 ± 0.28 | (*n* = 8)<br>0.92 ± 0.59 | (*n* = 6)<br>1.25 ± 0.87 | (*n* = 8)<br>1.43 ± 1.27 | NS | Age*Diet (NS, p = 0.077) | NS | NS | Age*Diet (NS, 3.25) | NS | 7 | 56 | 63 |
| ***Atp6v1g1*** | (*n* = 8)<br>1.07 ± 0.43 | (*n* = 10)<br>1.12 ± 0.45 | (*n* = 8)<br>1.00 ± 0.36 | (*n* = 7)<br>1.01 ± 0.54 | (*n* = 9)<br>1.26 ± 0.82 | (*n* = 8)<br>1.39 ± 0.51 | (*n* = 6)<br>1.04 ± 0.37 | (*n* = 8)<br>1.16 ± 0.55 | **Glucose AUC (p < 0.001)**<br>Age (NS, 0.086)<br>Diet (p = 0.014)<br>Sex (p = 0.015) | NS | NS | **Glucose AUC (15.41)**<br>Age (NS, 3.07)<br>Diet (6.51)<br>Sex (6.33) | NS | NS | 8** | 53 | 61 |

**p- and F-values are reported for p < 0.10. Statistical significance is defined as p < 0.05. NS = No significance. Bolded Glucose AUC indicates a significant covariate with adjusted p-values.**

***Degrees of freedom for 3-way ANOVA (1 for each of 3 factors, 1 for each 2-way interaction between 3 factors, 1 for a 3-way interaction)**

****Degrees of freedom for 3-way ANCOVA (1 for a covariate, 1 for each of 3 factors, 1 for each 2-way interaction between 3 factors, 1 for a 3-way interaction)**

**LFD = Low-fat Diet HFD = High-fat Diet**

**Aged HFD Females had smaller n because 2 mice were excluded from all analyses due to tumors after diet intervention**
**Aged LFD Females had smaller n because 1 mouse died before the end of the diet intervention**
**Aged LFD Males had smaller n because 1 mouse died before the end of the diet intervention**

**Table S10.** Fluorescent advanced glycation end products (fAGEs) results from flushed left humeri. Presented as mean ± SD.

| | Female | | | | Male | | | | Main Effects | 2-way Interactions | 3-way Interaction | Main Effect F-values | 2-way Interactions F-values | 3-way Interaction F-values | Degrees of Freedom* | Error | Total |
|---|---|---|---|---|---|---|---|---|---|---|---|---|---|---|---|---|---|
| | Young (5-month) | | Aging (22-month) | | Young (5-month) | | Aging (22-month) | | | | | | | | | | |
| | LFD | HFD | LFD | HFD | LFD | HFD | LFD | HFD | | | | | | | | | |
| | (*n = 10*) | (*n = 9*) | (*n = 9*) | (*n = 6*) | (*n = 10*) | (*n = 10*) | (*n = 6*) | (*n = 7*) | | | | | | | | | |
| **fAGEs (ng Q / mg Collagen)** | 351 ± 173 | 340 ± 87 | 296 ± 95 | 212 ± 68 | 315 ± 162 | 256 ± 65 | 275 ± 89 | 393 ± 131 | NS | NS | Sex*Age*Diet (p = 0.033) | NS | NS | Sex*Age*Diet (4.76) | 7 | 59 | 66 |

**p- and F-values are reported for p < 0.10. Statistical significance is defined as p < 0.05. NS = No significance**

***Degrees of freedom for 3-way ANOVA (1 for each of 3 factors, 1 for each 2-way interaction between 3 factors, 1 for a 3-way interaction)**

**LFD = Low-fat Diet** **HFD = High-fat Diet**

**Aged HFD Females had smaller n because 2 mice were excluded from all analyses due to tumors after diet intervention**
**Aged LFD Females had smaller n because 1 mouse died before the end of the diet intervention**
**Aged LFD Males had smaller n because 1 mouse died before the end of the diet intervention**

**Table S11.** Hydrated Raman spectroscopy was performed on the distal halves of left femora. Below are results from the periosteal surface and endocortical region (0-30% cortical thickness). The endocortical region presents measures describing the median values in the region, the gradient of the measure in the region (slope of linear fit of the data) and the variance of the data as calculated by the root mean square deviation (RMSD from residuals of linear fit). Presented as mean ± SD.

| | Female | | | | Male | | | | Main Effects | 2-way Interactions | 3-way Interaction | Main Effect F-values | 2-way Interactions F-values | 3-way Interaction F-values | Degrees of Freedom* | Error | Total |
|---|---|---|---|---|---|---|---|---|---|---|---|---|---|---|---|---|---|
| | Young (5-month) | | Aging (22-month) | | Young (5-month) | | Aging (22-month) | | | | | | | | | | |
| | LFD | HFD | LFD | HFD | LFD | HFD | LFD | HFD | | | | | | | | | |
| **Periosteal** | | | | | | | | | | | | | | | | | |
| **P. Median $v_2PO_4$:Amide III (Mineral: Matrix)** | *(n = 4)*<br>3.67 ± 0.33 | *(n = 4)*<br>3.51 ± 0.35 | *(n = 3)*<br>3.62 ± 0.67 | *(n = 3)*<br>3.76 ± 0.44 | *(n = 10)*<br>3.24 ± 0.28 | *(n = 10)*<br>3.34 ± 0.33 | *(n = 8)*<br>3.19 ± 0.20 | *(n = 8)*<br>3.02 ± 0.37 | Age (NS)<br>Diet (NS)<br>Sex ($p < 0.001$) | NS | NS | Age (NS)<br>Diet (NS)<br>Sex (16.48) | NS | NS | 7 | 42 | 49 |
| **P. Median $v_1CO_3$:$v_1PO_4$ (Carbonate:Phosphate)** | *(n = 4)*<br>0.111 ± 0.008 | *(n = 4)*<br>0.104 ± 0.003 | *(n = 3)*<br>0.116 ± 0.011 | *(n = 3)*<br>0.127 ± 0.007 | *(n = 10)*<br>0.109 ± 0.006 | *(n = 10)*<br>0.113 ± 0.010 | *(n = 8)*<br>0.122 ± 0.012 | *(n = 8)*<br>0.126 ± 0.014 | Age ($p < 0.001$)<br>Diet (NS)<br>Sex (NS) | NS | Sex*Age*Diet (NS, p = 0.096) | Age (21.02)<br>Diet (NS)<br>Sex (NS) | NS | Sex*Age*Diet (NS, 2.91) | 7 | 42 | 49 |
| **P. Median 1/ FWHM $v_1PO_4$ (Crystallinity)** | *(n = 4)*<br>0.0511 ± 0.0015 | *(n = 4)*<br>0.0517 ± 0.0012 | *(n = 3)*<br>0.0498 ± 0.0008 | *(n = 3)*<br>0.0506 ± 0.0013 | *(n = 10)*<br>0.0506 ± 0.0011 | *(n = 10)*<br>0.0505 ± 0.0020 | *(n = 8)*<br>0.0517 ± 0.0013 | *(n = 8)*<br>0.0519 ± 0.0016 | NS | Sex*Age (p = 0.016) | NS | NS | Sex*Age (6.26) | NS | 7 | 42 | 49 |
| **P. Median 1670/1640 $cm^{-1}$ (Matrix Order)** | *(n = 4)*<br>0.829 ± 0.029 | *(n = 4)*<br>0.861 ± 0.022 | *(n = 3)*<br>0.837 ± 0.061 | *(n = 3)*<br>0.783 ± 0.036 | *(n = 10)*<br>0.839 ± 0.056 | *(n = 10)*<br>0.829 ±0.065 | *(n = 8)*<br>0.838 ± 0.024 | *(n = 8)*<br>0.852 ± 0.034 | NS | NS | Sex*Age*Diet (NS, p = 0.072) | NS | NS | Sex*Age*Diet (NS, 3.41) | 7 | 42 | 49 |
| **P. Median 1670/1610 $cm^{-1}$ (Matrix Order)** | *(n = 4)*<br>0.824 ± 0.032 | *(n = 4)*<br>0.873 ± 0.038 | *(n = 3)*<br>0.835 ± 0.105 | *(n = 3)*<br>0.763 ± 0.056 | *(n = 10)*<br>0.841 ± 0.079 | *(n = 10)*<br>0.837 ± 0.101 | *(n = 8)*<br>0.819 ± 0.042 | *(n = 8)*<br>0.866 ± 0.066 | NS | NS | Sex*Age*Diet (NS, p = 0.073) | NS | NS | Sex*Age*Diet (NS, 3.38) | 7 | 42 | 49 |
| **Endocortical (0-30% Cortical Thickness)** | | | | | | | | | | | | | | | | | |
| **E. Median $v_2PO_4$:Amide III (Mineral: Matrix)** | *(n = 4)*<br>3.85 ± 0.22 | *(n = 4)*<br>4.49 ± 0.22 | *(n = 3)*<br>4.06 ± 0.25 | *(n = 4)*<br>3.97 ± 0.38 | *(n = 10)*<br>3.55 ± 0.33 | *(n = 10)*<br>3.59 ± 0.25 | *(n = 7)*<br>3.70 ± 0.28 | *(n = 7)*<br>3.74 ± 0.20 | NS | NS | Sex*Age*Diet (p = 0.039) | NS | NS | Sex*Age*Diet (4.57) | 7 | 41 | 48 |
| **E. Median $v_1CO_3$:$v_1PO_4$ (Carbonate:Phosphate)** | *(n = 4)*<br>0.103 ± 0.003 | *(n = 4)*<br>0.093 ± 0.014 | *(n = 3)*<br>0.105 ± 0.006 | *(n = 4)*<br>0.094 ± 0.014 | *(n = 10)*<br>0.108 ± 0.009 | *(n = 10)*<br>0.110 ± 0.010 | *(n = 8)*<br>0.113 ± 0.011 | *(n = 8)*<br>0.113 ± 0.013 | NS | **Glucose AUC (p = 0.023)**<br>Sex*Diet (p = 0.020) | NS | NS | **Glucose AUC (5.54)**<br>Sex*Diet (5.85) | NS | 8** | 41 | 49 |
| **E. Median 1/ FWHM $v_1PO_4$ (Crystallinity)** | *(n = 4)*<br>0.0528 ± 0.0024 | *(n = 4)*<br>0.0420 ± 0.0073 | *(n = 3)*<br>0.0496 ± 0.0066 | *(n = 4)*<br>0.0450 ± 0.0087 | *(n = 10)*<br>0.0514 ± 0.0046 | *(n = 10)*<br>0.0520 ± 0.0036 | *(n = 8)*<br>0.0516 ± 0.0046 | *(n = 8)*<br>0.0512 ± 0.0040 | NS | Sex*Diet (p = 0.045) | NS | NS | Sex*Diet (4.26) | NS | 7 | 43 | 50 |
| **E. Median 1670/1640 $cm^{-1}$ (Matrix Order)** | *(n = 4)*<br>1.19 ± 0.17 | *(n = 4)*<br>1.11 ± 0.09 | *(n = 3)*<br>1.22 ± 0.11 | *(n = 4)*<br>1.15 ± 0.21 | *(n = 10)*<br>1.11 ± 0.12 | *(n = 10)*<br>1.15 ± 0.12 | *(n = 8)*<br>1.19 ± 0.08 | *(n = 8)*<br>1.15 ± 0.12 | NS | NS | NS | NS | NS | NS | 7 | 43 | 50 |
| **E. Median 1670/1610 $cm^{-1}$ (Matrix Order)** | *(n = 4)*<br>3.29 ± 1.52 | *(n = 4)*<br>1.92 ± 0.61 | *(n = 3)*<br>3.58 ± 1.45 | *(n = 4)*<br>3.02 ± 1.70 | *(n = 10)*<br>2.33 ± 0.78 | *(n = 10)*<br>2.55 ± 0.72 | *(n = 8)*<br>2.81 ± 0.85 | *(n = 8)*<br>2.65 ± 0.91 | NS | NS | NS | NS | NS | NS | 7 | 43 | 50 |
| **E. Gradient $v_2PO_4$:Amide III (Slope Mineral: Matrix)** | *(n = 4)*<br>8.30E-3 ± 26.6E-3 | *(n = 4)*<br>7.32E-3 ± 14.9E-3 | *(n = 3)*<br>21.7E-3 ± 31.8E-3 | *(n = 4)*<br>2.42E-3 ± 15.0E-3 | *(n = 10)*<br>9.63E-3 ± 19.8E-3 | *(n = 10)*<br>17.2E-3 ± 19.5E-3 | *(n = 8)*<br>37.6E-3 ± 22.0E-3 | *(n = 8)*<br>7.60E-3 ± 47.6E-3 | **Glucose AUC (p = 0.006)**<br>Age (NS)<br>Diet (NS)<br>Sex (NS, p = 0.096) | NS | NS | **Glucose AUC (8.56)**<br>Age (NS)<br>Diet (NS)<br>Sex (NS, 2.91) | NS | NS | 8** | 41 | 49 |
| **E. Gradient $v_1CO_3$:$v_1PO_4$ (Slope Carbonate:Phosphate)** | *(n = 4)*<br>-2.02E-4 ± 3.10E-4 | *(n = 4)*<br>4.72E-4 ± 3.26E-4 | *(n = 3)*<br>0.87E-4 ± 2.13E-4 | *(n = 4)*<br>-1.39E-4 ± 3.95E-4 | *(n = 10)*<br>-2.37E-4 ± 6.63E-4 | *(n = 9)*<br>1.92E-4 ± 3.10E-4 | *(n = 8)*<br>-1.00E-4 ± 5.41E-4 | *(n = 8)*<br>-0.45E-4 ± 3.87E-4 | NS | Age*Diet (p = 0.032) | NS | NS | Age*Diet (4.92) | NS | 7 | 42 | 49 |
| **E. Gradient 1/ FWHM $v_1PO_4$ (Slope Crystallinity)** | *(n = 4)*<br>-0.30E-5 ± 2.5E-5 | *(n = 4)*<br>11.4E-5 ± 13.1E-5 | *(n = 3)*<br>2.10E-5 ± 4.30E-5 | *(n = 4)*<br>1.60E-5 ± 3.50E-5 | *(n = 10)*<br>1.20E-5 ± 3.90E-5 | *(n = 9)*<br>0.70E-5 ± 5.50E-5 | *(n = 7)*<br>-4.90E-5 ± 4.2E-5 | *(n = 8)*<br>-1.00E-5 ± 8.6E-5 | NS | NS | Sex*Age*Diet (p = 0.040) | NS | NS | Sex*Age*Diet (4.50) | 7 | 41 | 48 |
| **E. Gradient 1670/1640 $cm^{-1}$ (Slope Matrix Order)** | *(n = 4)*<br>0.01E-3 ± 2.62E-3 | *(n = 4)*<br>3.53E-3 ± 2.60E-3 | *(n = 3)*<br>1.60E-3 ± 6.74E-3 | *(n = 4)*<br>3.20E-3 ± 3.33E-3 | *(n = 10)*<br>3.04E-3 ± 5.35E-3 | *(n = 10)*<br>3.55E-3 ± 3.82E-3 | *(n = 8)*<br>4.98E-3 ± 6.43E-3 | *(n = 8)*<br>1.37E-3 ± 5.53E-3 | NS | NS | NS | NS | NS | NS | 7 | 43 | 50 |
| **E. Gradient 1670/1610 $cm^{-1}$ (Slope Matrix Order)** | *(n = 4)*<br>0.110 ± 0.281 | *(n = 4)*<br>0.018 ± 0.008 | *(n = 3)*<br>-0.005 ± 0.152 | *(n = 3)*<br>0.015 ± 0.004 | *(n = 10)*<br>0.029 ± 0.045 | *(n = 10)*<br>0.028 ± 0.024 | *(n = 8)*<br>0.035 ± 0.043 | *(n = 8)*<br>0.025 ± 0.053 | NS | NS | NS | NS | NS | NS | 7 | 42 | 49 |
| **E. Heterogeneity $v_2PO_4$:Amide III (RMSD Mineral: Matrix)** | *(n = 4)*<br>0.437 ± 0.195 | *(n = 4)*<br>0.436 ± 0.122 | *(n = 3)*<br>0.371 ± 0.130 | *(n = 4)*<br>0.490 ± 0.213 | *(n = 10)*<br>0.487 ± 0.208 | *(n = 10)*<br>0.433 ± 0.190 | *(n = 8)*<br>0.608 ± 0.359 | *(n = 8)*<br>0.541 ± 0.227 | NS | NS | NS | NS | NS | NS | 7 | 43 | 50 |
| **E. Heterogeneity $v_1CO_3$:$v_1PO_4$ (RMSD Carbonate:Phosphate)** | *(n = 4)*<br>8.36E-3 ± 5.62E-3 | *(n = 4)*<br>6.35E-3 ± 2.12E-3 | *(n = 3)*<br>6.51E-3 ± 3.17E-3 | *(n = 4)*<br>6.16E-3 ± 3.05E-3 | *(n = 10)*<br>11.7E-3 ± 6.1E-3 | *(n = 9)*<br>8.08E-3 ± 3.86E-3 | *(n = 8)*<br>8.17E-3 ± 4.63E-3 | *(n = 8)*<br>10.2E-3 ± 4.6E-3 | Sex (p = 0.033) | NS | NS | Sex (4.89) | NS | NS | 7 | 42 | 49 |
| **E. Heterogeneity 1/ FWHM $v_1PO_4$ (RMSD Crystallinity)** | *(n = 4)*<br>1.41E-3 ± 0.46E-3 | *(n = 4)*<br>1.13E-3 ± 0.27E-3 | *(n = 3)*<br>1.22E-3 ± 0.42E-3 | *(n = 4)*<br>1.11E-3 ± 0.27E-3 | *(n = 10)*<br>1.18E-3 ± 0.66E-3 | *(n = 10)*<br>1.57E-3 ± 0.74E-3 | *(n = 7)*<br>1.69E-3 ± 0.61E-3 | *(n = 7)*<br>1.33E-3 ± 0.31E-3 | NS | NS | NS | NS | NS | NS | 7 | 41 | 48 |
| **E. Heterogeneity 1670/1640 $cm^{-1}$ (RMSD Matrix Order)** | *(n = 4)*<br>0.0756 ± 0.0332 | *(n = 4)*<br>0.0578 ± 0.0277 | *(n = 3)*<br>0.0554 ± 0.0058 | *(n = 4)*<br>0.0606 ± 0.0295 | *(n = 10)*<br>0.0517 ± 0.0144 | *(n = 10)*<br>0.0560 ± 0.0224 | *(n = 7)*<br>0.0529 ± 0.0092 | *(n = 8)*<br>0.0695 ± 0.0233 | NS | NS | NS | NS | NS | NS | 7 | 42 | 49 |
| **E. Heterogeneity 1670/1610 $cm^{-1}$ (RMSD Matrix Order)** | *(n = 4)*<br>3.04 ± 3.00 | *(n = 4)*<br>0.316 ± 0.255 | *(n = 3)*<br>3.42 ± 1.93 | *(n = 4)*<br>1.51 ± 2.07 | *(n = 10)*<br>0.632 ± 0.537 | *(n = 9)*<br>0.446 ± 0.182 | *(n = 8)*<br>0.603 ± 0.222 | *(n = 7)*<br>0.781 ± 0.556 | Age (NS, p = 0.098) | Sex*Diet (p = 0.001) | NS | Age (NS, 2.87) | Sex*Diet (12.94) | NS | 7 | 41 | 48 |

**p- and F-values are reported for $p < 0.10$. Statistical significance is defined as $p < 0.05$. NS = No significance. Bolded Glucose AUC indicates a significant covariate with adjusted p-values.**
***Degrees of freedom for 3-way ANOVA (1 for each of 3 factors, 1 for each 2-way interaction between 3 factors, 1 for a 3-way interaction)**
****Degrees of freedom for 3-way ANCOVA (1 for a covariate, 1 for each of 3 factors, 1 for each 2-way interaction between 3 factors, 1 for a 3-way interaction)**

**LFD = Low-fat Diet** **HFD = High-fat Diet**

**All female groups have smaller n due to sample availability (1/2 unavailable)**

**Table S12.**Summary of primers used for qRT-PCR on flushed tibiae bone.

| Gene | Forward Primer | Reverse Primer |
|---|---|---|
| | **Bone** | |
| ***Tnfrsf11b (Opg)*** | AGAGCAAACCTTCCAGCTGC | CTGCTCTGTGGTGAGGTTCG |
| ***Tnfsf11 (Rankl)*** | CCAAGATCTCTAACATGACG | CACCATCAGCTGAAGATAGT |
| ***Atp6v1g1*** | CCGTTCTCTCAGCCCAAAGT | CTCCGGTTCTTTCGCTTGC |
| ***Atp6v0d2*** | TCTTGAGTTTGAGGCCGACAG | GCAACCCCTCTGGATAGAGC |
| ***Acp5*** | CGTCTCTGCACAGATTGCAT | AAGCGCAAACGGTAGTAAGG |
| ***Mmp2*** | AACGGTCGGGAATACAGCAG | GTAAACAAGGCTTCATGGGG |
| ***Mmp13*** | CGGGAATCCTGAAGAAGTCTACA | CTAAGCCAAAGAAAGATTGCATTTC |
| ***Mmp14*** | AGGAGACGGAGGTGATCATCATTG | GTCCCATGGCGTCTGAAGA |
| ***Ctsk*** | GAGGGCCAACTCAAGAAGAA | GCCGTGGCGTTATACATACA |
| | **Housekeeper** | |
| ***Tbp*** | ACCTTATGCTCAGGGCTTGG | GCCATAAGGCATCATTGGAC |
| ***B2m*** | ACAGTTCCACCCGCCTCACATT | TAGAAAGACCAGTCCTTGCTGAAG |
| ***Hsp90ab1*** | CCTGAAGGTCATCCGCAAGAAC | GGCGTCGGTTAGTGGAATCTTC |
| ***Gapdh*** | TCCCACTCTTCCACCTTCGA | AGTTGGGATAGGGCCTCTCTTG |

**Table S13.** Summary of measures that have different conclusions if the outliers are not excluded from analysis.

| Measure | Without Outliers (see other tables) | With Outliers | Different Conclusion(s) with Outliers Included | Outlier Information (Grubb's Test) | Outlier ID |
|---|---|---|---|---|---|
| **Epididymal Adipose Depot Mass (g)** | **Glucose AUC (p < 0.001)**<br>Sex*Age (p < 0.001)<br>Sex*Diet (p < 0.001)<br>Age*Diet (0.037)<br>See Table S1 | **Glucose AUC (p = 0.003)**<br>Sex*Age (p < 0.001)<br>Sex*Diet (p = 0.014)<br>Age*Diet (NS) | Sex*Age and Sex*Diet interactions are the same<br>**No** Age*Diet interactions (i.e. young HFD mice have greater adipose tissue compared to young LFD) | Young HFD female (+50% from nearest in group)<br>Aged HFD male (+170% from nearest in group) | AD26; AD43 |
| **IL-6 (pg/mL)** | Sex (NS)<br>Age (p < 0.001)<br>Diet (0.047)<br>See Table S2 | Sex (NS)<br>Age (p < 0.001)<br>Diet (NS) | Same age conclusions<br>**No** diet effect (HFD reduces IL-6 compared to LFD mice) | Young HFD female (+5000% from nearest in group)<br>Young and Aged HFD males (+110% and +475 from nearest in their respective groups) | AD30; AD41; AD56 |
| **IL-10 (pg/mL)** | Sex*Age (p = 0.036)<br>See Table S2 | Sex (p = 0.004)<br>Age (p < 0.001) | Male -36% compared to female mice<br>Aged +107% compared to young mice | Young HFD female (+115% from nearest in group)<br>Aged LFD and HFD males (+440% and +250% from nearest in their respective groups) | AD30; AD45; AD68 |
| **FGF21 (ng/mL)** | Sex (p = 0.004)<br>Age (NS, p = 0.066)<br>See Table S2 | Sex (p = 0.004)<br>Age (p = 0.039) | Sex main effect is the same<br>Aged +38% compared to young mice | Aged LFD female (+50% from nearest in group) | AD16 |
| **Ct. Ar (mm²)** | Diet (NS, 0.052)<br>Sex*Age (p = 0.028)<br>See Table S3 | Diet (p = 0.033)<br>Sex*Age (p = 0.048) | Sex*Age conclusions are the same<br>HFD -4% compared to LFD mice | Young HFD female (-12% from nearest in group) | AD1 |
| **Ct. Th (mm)** | Sex (p < 0.001)<br>Age (p < 0.001)<br>See Table S3 | Sex*Age (p = 0.036) | Aged mice of both sexes have lower Ct. Th. (the same)<br>There is a sex difference in young mice (young males -8% compared to young females) but not aged mice | Young LFD and HFD females (-8% and -27% from nearest in their respective groups)<br>Age LFD female (-14% from nearest in group) | AD1; AD10; AD18 |
| **Imin (mm⁴)** | Sex (NS, p = 0.072)<br>Age (p < 0.001)<br>See Table S3 | Sex (p = 0.043)<br>Age (p < 0.001) | Male +6% compared to female mice<br>Same age conclusions | Young LFD male (+33% from nearest in group) | AD54 |
| ***Atp6v1g1*** | **Glucose AUC (p < 0.001)**<br>Sex (p =0.015)<br>Diet (p = 0.014)<br>See Table S9 | **Glucose AUC (p = 0.001)**<br>Sex*Diet (p = 0.049) | Nothing significant after correction | Young and aged LFD females (+76 % and +16000% from nearest in their respective groups) | AD8; AD15 |
| ***Acp5*** | Sex (p = 0.015)<br>Age (p = 0.036)<br>See Table S9 | Sex (NS, p = 0.052)<br>Age (p = 0.025) | **No** sex effect (i.e. male -36% compared to female mice)<br>Same age conclusions | Aged LFD and HFD females (+16000% and +170% from nearest in their respective groups)<br>Aged LFD male (+100% from nearest in group) | AD11; AD15; AD49 |
| ***Mmp13*** | Sex*Diet (p = 0.047)<br>See Table S9 | Sex (p = 0.028) | Male -39% compared to female mice | Aged LFD and HFD females (+14000% and +115% from nearest in their respective groups) | AD11; AD15 |
| **E. Median $v_2PO_4$:Amide III (Mineral: Matrix)** | Sex*Age*Diet (p = 0.039)<br>See Table S11 | Sex (p = 0.002) | Male -10% compared to female mice | Aged LFD and HFD male mice (both are +30% from nearest in their respective groups) – Both have high chance of error due to fewer points from plastic contamination | AD45; AD68 |

**Bolded Glucose AUC indicates a significant covariate with adjust p-values**

**LFD = Low-fat Diet** **HFD = High-fat Diet**
